# Structural characterization of an extracellular contractile injection system from *Photorhabdus luminescens* in extended and contracted states

**DOI:** 10.1101/2025.04.20.649488

**Authors:** Leyre Marín-Arraiza, Aritz Roa-Eguiara, Tillmann Pape, Nicholas Sofos, Ivo Alexander Hendriks, Michael Lund Nielsen, Eva Maria Steiner-Rebrova, Nicholas M. I. Taylor

## Abstract

Contractile injection systems (CISs) are phage-tail-like nanosyringes that mediate bacterial interactions by puncturing target cell membranes. Within these systems, Photorhabdus Virulence Cassettes (PVCs) can translocate toxins across eukaryotic target cell membranes. The structure of a PVC has been described at atomic level and engineered to deliver diverse protein cargoes into non-natively-targeted organisms. Despite the structural insights into several CISs, information on PVCs from other species and details on the contraction mechanism remain limited. Here, we present the single-particle cryo-electron microscopy structure of *Pl*PVC1, a PVC from the nematode symbiont and insect pathogen *Photorhabdus luminescens DJC*, in both extended and contracted states. Our structure displays distinct structural features that differ from other CISs, such as a cage surrounding the central spike, a larger sheath adaptor, and a plug exposed to the tube lumen. Moreover, we present the structures of the *Pl*PVC1 fiber as well as the baseplate of the contracted particle, yielding insight into the contraction mechanism. This study provides structural details of the contracted state of the *Pl*PVC1 particle and supports the model in which contraction is triggered. Furthermore, it facilitates the comparison of *Pl*PVC1 with other contractile systems and expands the scope of engineering opportunities for future biomedical and biotechnological applications.

## INTRODUCTION

Microbial communities coexist and compete in their natural environments by interacting with their surroundings and other organisms^1^. These interactions often involve the translocation of macromolecules across cell membranes, mediating processes such as cellular communication and defense^2,3^. To facilitate this, many bacterial species have evolved a plethora of specialized machineries, the contractile injection systems (CISs), macromolecular nanosyringes distinct from one another but evolutionarily related to bacteriophage tails^4^. These particles share a common structure consisting of a contractile sheath wrapped around a rigid tube, sharpened with a central spike, an assembly used for payload delivery^4^. The central spike is surrounded by a baseplate complex equipped with fibers for host recognition. The baseplate triggers contraction upon specific sensing of the target cell through the fiber network^5^. Contraction of the sheath drives the spiked tube outward, piercing the target cell membrane and injecting the payload.

Based on the anchoring mechanism of the baseplate to the membrane prior to action, bacterial CISs are commonly classified into type VI secretion systems (T6SSs) and extracellular contractile injection systems (eCISs). T6SSs are cell-wall anchored CISs, widespread among Gram-negative bacteria^6^, which deliver bacterial effectors into target prokaryotic or eukaryotic cells by being pushed out of the bacterial membrane^7,8^. Differently, eCISs attack target cells from the extracellular environment. Several eCISs have been studied, each produced by different microorganisms and exhibiting distinct structural and functional features. Tail-like bacteriocins, or tailocins, are a broad family of eCISs^9^, with the most well-studied example being the R-pyocin encoded by *Pseudomonas aeruginosa*^10,11^. R-pyocins bind to the receptor sites on the lipopolysaccharide of the target cell^12^, and disrupt the membrane potential after puncturing of the cell membrane^13^. Metamorphosis-associated contractile structures (MACs), produced by *Pseudoalteromonas luteoviolacea*, arrange in ordered bundles^14^ and carry effectors that are necessary to induce metamorphosis^15^ and kill eukaryotic cells^16^. The well-characterized antifeeding prophage (AFP), produced by *Serratia entomophila*, acts as a delivery vehicle for the insecticidal toxin Afp18 and causes amber disease in the New Zealand grass grub^17–19^. Recently, a bacterial CIS found in *Algoriphagus machipongonensis* (AlgoCIS) exhibits structural differences compared to canonical contractile systems, presenting a cap adaptor, a plug harbored inside the tube lumen, and a cage-like structure around the spike^20^.

Photorhabdus Virulence Cassettes (PVCs) are eCISs produced by bacteria of the *Photorhabdus* genus, which can translocate toxins across eukaryotic target membranes^21,22^. Different *Photorhabdus* species encode distinct copies of *pvc* operons in their genome, each associated with unique putative effector genes^23,24^. Notably, PVC-like genes are broadly distributed in the genomes of both prokaryotes and archaea^25,26^, suggesting that these systems may represent an ancient mechanism contributing to microbial evolution and functional specialization^23^. The cryo-electron microscopy (cryo-EM) structure of PVCpnf, a PVC from *P. asymbiotica*, features a contractile phage-tail-like particle^27^, and its target specificity is mediated by the recognition of cellular receptors by the tail fibers. These fibers can be genetically engineered to retarget the particle against non-natively-targeted organisms with high efficiency^28^. Moreover, PVCpnf is being studied for its potential to load diverse protein cargoes, which could be delivered to different targeted cells, both *in vitro* and *in vivo*^28,29^. This programmable capability provides the opportunity to customize PVCs for specific therapeutic applications.

Currently, structural and functional studies of PVCs are limited to PVCpnf from *P. asymbiotica*, and no cryo-EM structures have been reported for PVCs from other *Photorhabdus* species. Given that *P. luminescens* is a nematode symbiont and insect pathogen^30^ that encodes six *pvc* operons in its genome^23,24^, the characterization of PVCs from this species could significantly contribute to our understanding of the potential role of eCISs in symbiosis and infection.

In this study, we use cryo-EM to characterize the high-resolution structure of a novel PVC particle from *Photorhabdus luminescens DJC* (*Pl*PVC1), in both its extended and contracted states, providing critical insights into its architecture and function. This system resembles other phage-tail-like particles, but with distinct structural features, such as the presence of a cage surrounding the central spike, a larger sheath adaptor, and a plug exposed to the tube lumen. Furthermore, the structures of the fiber and the baseplate of the contracted *Pl*PVC1 particle are solved, providing a comprehensive framework for understanding the contraction mechanism. The detailed structural characterization and comparison of the extended and contracted states support the model in which contraction is triggered upon target cell recognition by the fibers. These findings significantly advance our understanding of PVCs and contribute to their promising customization as biomedical tools, from biocontrol to precision therapy.

## RESULTS

### Overall structure of the *Pl*PVC1 particle

The *pvc* operon 1 from *Photorhabdus luminescens DJC* was cloned for expression in *Escherichia coli*. It comprises 16 open reading frames encoding the proteins conforming the *Pl*PVC1 particle (Pvc1 to Pvc16) (**Fig.1a**, **Sup.Fig.1**, **Sup.Tab.1**). Mass spectrometry (MS) confirmed the presence of all proteins in the purified sample. After expression, fully assembled *Pl*PVC1 particles were visualized with negative-staining electron microscopy (NS-EM) (**Sup.Fig.2a**). The length of purified *Pl*PVC1 particles in extended state was heterogeneous, with an average particle length of ∼280 nm (**Sup.Fig.2b**, **Sup.Tab.2**). Single-particle cryo-EM was used to determine the high-resolution structure of the *Pl*PVC1 particle components (**Sup.Figs.3-4**, **Sup.Tab.3**). The structures show similar features to other phage-tail-like particles^11,19,20,27^: a baseplate surrounded by tail fibers network and equipped with a central spike sharpened at the tip, a contractile trunk composed of sheath and inner tube, and a terminal cap at the apical end (**Fig.1b-1c**, **Sup.Fig.2a**). The *Pl*PVC1 particle generally follows 6-fold symmetry along its structure, with symmetry mismatches between the baseplate (6-fold), central spike (3-fold), and spike tip (1-fold). In the extended state, the outer diameter of the sheath reaches 162 Å, enclosing the inner tube, which has an outer and inner diameter of 80 Å and 45 Å, respectively. At the baseplate level, the particle diameter expands up to 280 Å (**Fig.1b**). Symmetry-based single-particle reconstruction was divided into regions – cap, baseplate, central spike, and fiber – and mask-based processing was used to improve the density resolution of specific parts (**Sup.Figs.3-4**, **Sup.Tab.3**).

**Figure 1.**
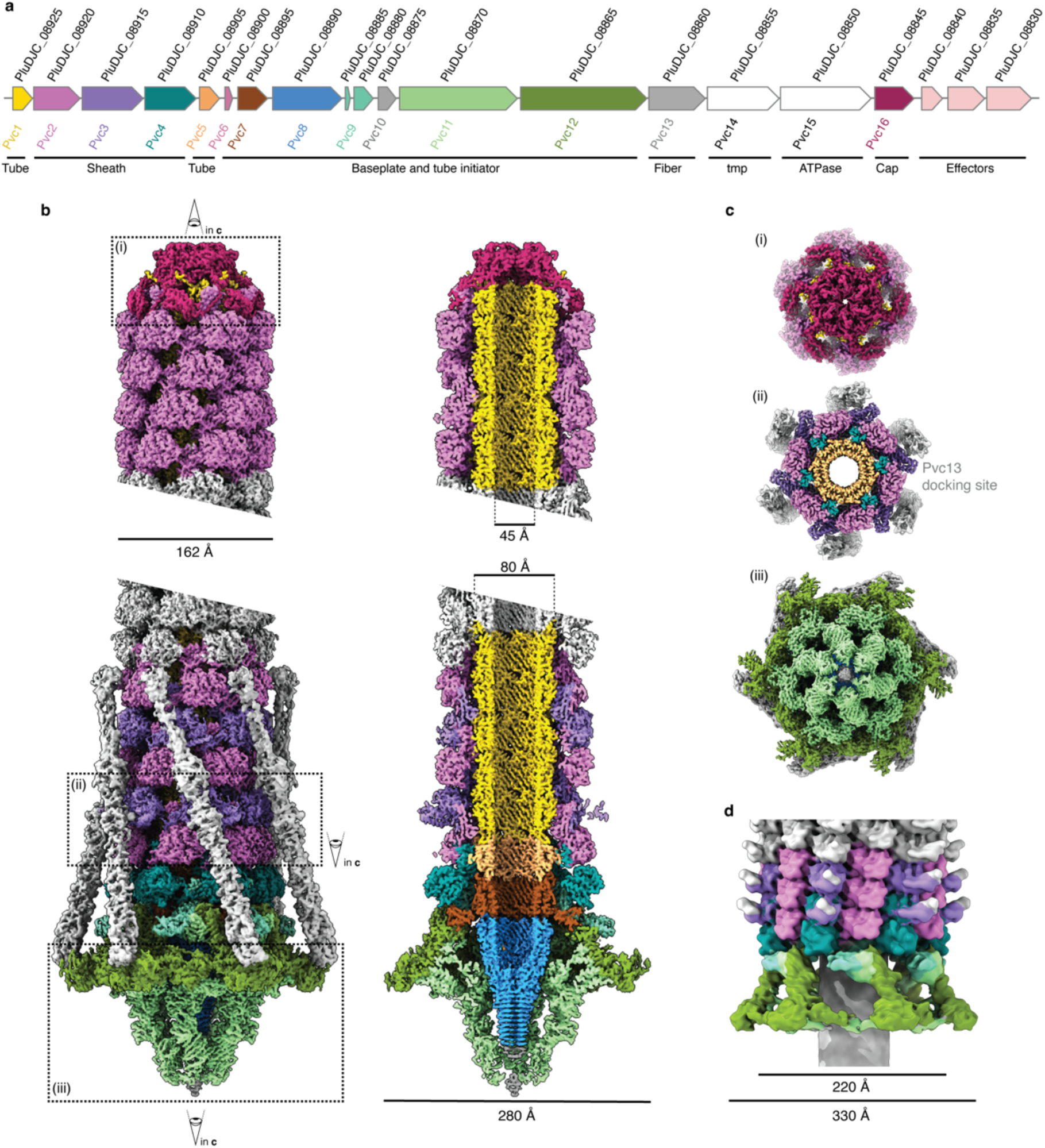
Overall cryo-EM structure of the *Pl*PVC1 particle in its extended and contracted states. **(a)** Schematic representation of the genomic organization of the *Pl-pvc1* cluster. The gene accession numbers are shown above the corresponding genes. **(b)** Cryo-EM maps of the *Pl*PVC1 particle in its extended state, in complete and slice views. The different structural subunits are colored according to **a**. **(c)** Horizontal cut-out views of the marked sections i-iii in **b**. (i) top view of the terminal cap; (ii) bottom view of the fiber docking site; (iii) bottom view of the baseplate. **(d)** Cryo-EM map of the *Pl*PVC1 particle in its contracted state, filtered to 10 Å. The different structural subunits are colored according to **a**.

The cap and baseplate density maps in the extended state were reconstructed to overall resolutions of 2.5 Å and 2.7 Å, respectively, applying 6-fold symmetry. The central spike was reconstructed to 2.8 Å by applying 3-fold symmetry. Using these cryo-EM maps, 14 proteins were located in the *Pl*PVC1 particle, with 12 having atomic models built. Proteins Pvc14 and Pvc15, which were present in the sample as verified by MS, were not located in any density map and thus were not modeled. The fibers were reconstructed individually, by local refinement in a symmetry-expanded particle set, to a resolution ranging from 4 Å to 6 Å. The AlphaFold model of a trimer of the fiber protein Pvc13 was fitted into the fiber density map, and the interaction between baseplate and fiber was built and refined (**Sup.Figs.3-4**, **Sup.Tab.3**).

Contraction of the *Pl*PVC1 particle was induced by exposing purified particles to 3 M urea^31,32^ (**Fig.1d**, **Sup.Fig.2c**). The contracted sheath was solved at 3.1 Å, and atomic models of the contracted sheath proteins were built. The baseplate of the contracted particle was solved at a resolution ranging from 4 Å to 10 Å, and atomic models of the baseplate proteins were rigid-body fitted in the density map (**Sup.Figs.3-4**, **Sup.Tab.3**).

### *Pl*PVC1 baseplate

The overall architecture of the *Pl*PVC1 baseplate in its extended state is similar to the one in PVCpnf^27^ and AFP^19^ particles and resembles a streamlined T4 inner baseplate^5^ (**Fig.2a**, **Sup.Fig.5**). The *Pl*PVC1 baseplate complex exhibits a 6-fold symmetrical assembly of the wedges (Pvc11 and Pvc12) surrounding the trimeric central spike (Pvc8), sharpened with the spike tip (Pvc10). The central spike prolongs from the inner tube, while the baseplate wedges are connected to the trunk of the particle through the sheath adaptor (Pvc9) (**Fig.2a**).

**Figure 2.**
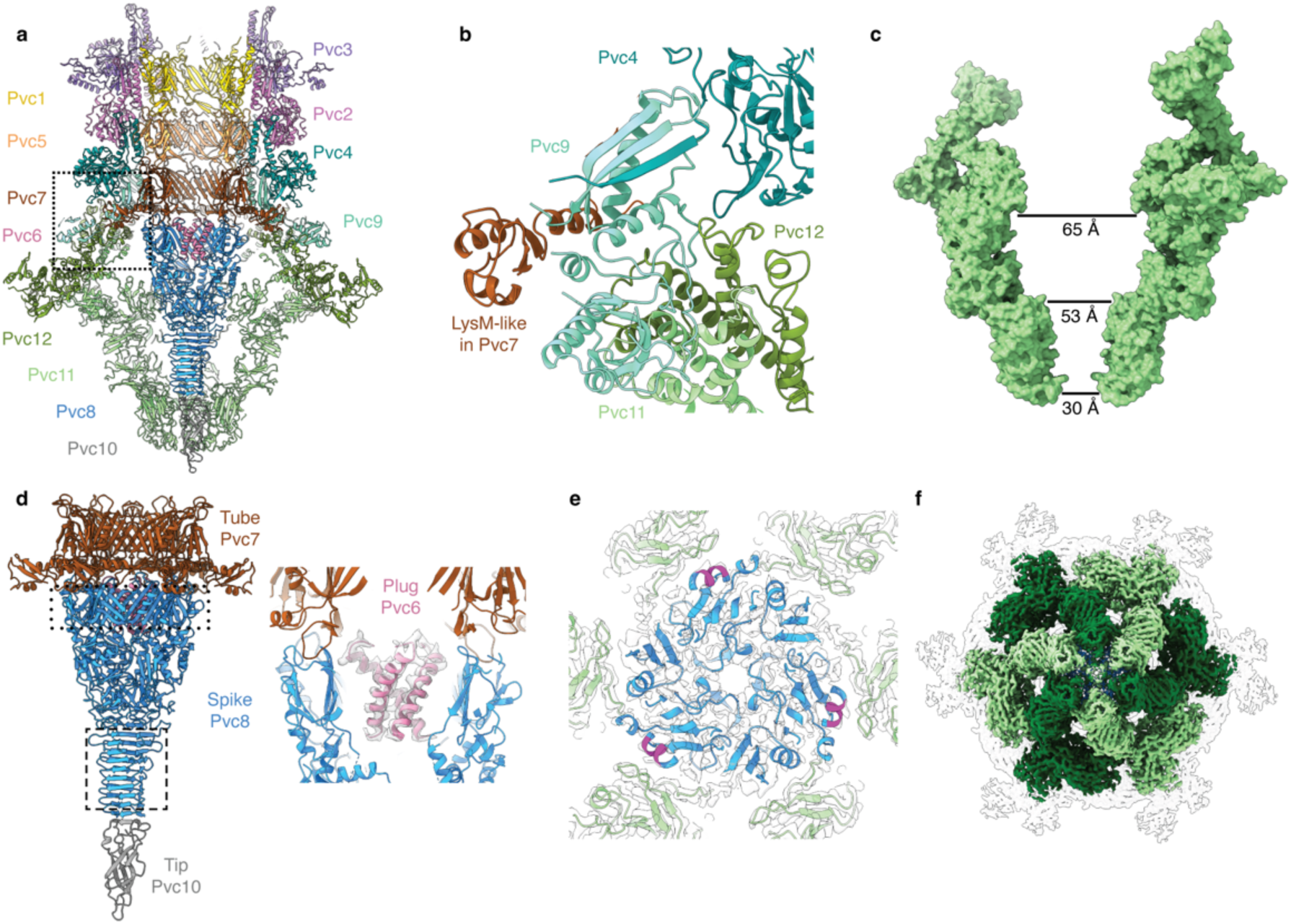
Organization and atomic model of the *Pl*PVC1 baseplate in its extended state. **(a)** Side cut-out view of the atomic model of the baseplate complex in the *Pl*PVC1 particle. The first layers of the sheath and tube are also shown. **(b)** Zoom-in view of the marked section in **a**, in ribbon diagram, showing interactions between the sheath adaptor Pvc9 and the sheath initiator Pvc4, LysM-like domain in Pvc7, and baseplate wedges Pvc11 and Pvc12. **(c)** Cut-out side view of surface diagram showing the spike cage in the *Pl*PVC1 baseplate. Measures between opposite Pvc11 subunits are labelled. **(d)** *Left:* Side view of the atomic model of the tube initiator, central spike, and spike tip in the *Pl*PVC1 particle. The dotted rectangle remarks the β-barrel conformation, transitioning between C6 and C3 symmetry. The dashed rectangle remarks the cone-shaped rigid spike. *Right*: Zoom-in view of the plug protein Pvc6 in the lumen of the inner tube, showing cryo-EM density and ribbon diagram. **(e)** Top cut-out view of the interactions between Pvc11 and the central spike Pvc8 in the upper part of the spike, depicted in magenta. The symmetry mismatch C6-C3 is circumvented by alternating interactions between subunits. **(f)** Horizontal bottom view of the configuration of the cage around the spike, with Pvc11 monomers colored according to their interaction with Pvc8; light green: contacting, dark green: non-contacting.

Pvc11 and Pvc12 arrange in heterodimers, similarly to the phage T4 [gp6]_2_-gp7 helical core bundle^5^. This [gp6]_2_-gp7-like core bundle in the baseplate wedges is conserved among other eCIS^11,19,20,27^, highlighting its importance in baseplate assembly and functionality. In the case of *Pl*PVC1, Pvc12 resembles a combination of T4 gp6 in the “gp6B” position and gp7, whereas Pvc11 is similar to T4 gp6 in the “gp6A” position (**Sup.Fig.6a**).

The sheath adaptor Pvc9 in *Pl*PVC1 is homologous to Pvc9 in PVCpnf, Afp9, Alg9, and gp25 in phage T4 (**Sup.Fig.6b**). Interestingly, Pvc9 is longer in the *Pl*PVC1 particle than in those other CISs and protrudes on top of Pvc12 and Pvc11 (**Fig.2b**, **Sup. Fig.7**, **Sup.Tabs.4-5**). As seen in other CISs^5,19,20,27^, the sheath adaptor acts as an interface between the baseplate wedges and the trunk of the particle, facilitating sheath orientation and assembly via several interactions between Pvc9 and baseplate, sheath, and tube proteins. Pvc9 strongly interacts with the sheath initiator Pvc4 and the tube initiator Pvc7, and docks on top of Pvc11 and Pvc12 (**Fig.2b**). Interaction-prediction analysis^33,34^ predicted Pvc9 interplay with proteins in the region where the sheath adaptor interacts with Pvc4, Pvc7, Pvc11, and Pvc12. Additionally, isolated interactions with carbohydrates were predicted for some residues in the protruded region, which is exposed toward the outer part of the particle (**Sup.Fig.6c**). This could lead to hypotheses of other roles for Pvc9, apart from organizing sheath orientation and assembly.

### *Pl*PVC1 baseplate cage

The baseplate of *Pl*PVC1 features an expanded cage around the central spike, formed by extensions of the protein Pvc11 (**Fig.2a,2c**). The cavity of the cage ranges between 65 Å and 30 Å in diameter (**Fig.2c**). This cage-like structure was not determined in PVCpnf^27^ or in AFP^19^, but was identified in the AlgoCIS particle^20^ (**Sup.Fig.5**, **Sup.Fig.6d, Sup. Fig.7**, **Sup.Tabs.4-5**). When compared to Alg11, both Pvc11 and Alg11 present an analogous fold in their N-terminal and C-terminal regions, which interact with Pvc12/Alg12. The folding of the cage extensions is also similar in both cases, although it is shorter in Pvc11 (**Sup.Fig.6d**). The inner surface of the Pvc11 cage is mainly negatively charged, in contrast to the surrounded spike, which presents a positively charged outer surface (**Sup.Fig.6e**). Structure-based bioinformatic analysis^35,36^ showed structural homology between carbohydrate-binding proteins and Pvc11 extensions, in consensus with similar results for Alg11^20^. In addition, interaction-prediction analysis^33,34^ predicted putative interactions with lipids and carbohydrates in the cage extensions (**Sup.Fig.6f**).

### *Pl*PVC1 central spike

The central spike in *Pl*PVC1 is composed of three copies of the protein Pvc8. It extends from the inner tube and is sharpened by one copy of the spike tip protein Pvc10 (**Fig.2a,2d**). There are three main interactions between Pvc8 and Pvc11, accommodating the association between central hub and baseplate wedges (**Fig.2e**). These interactions lead to the specific arrangement of Pvc11 around Pvc8, in an alternating pattern of contacting and non-contacting monomers, which is presumably important for baseplate assembly and stabilization (**Fig.2f**).

Similar to VgrG in T6SS^6^, Pvc8 is a fusion protein of the central hub genes from phage T4. The N-terminal region of Pvc8, which functions as the symmetry adaptor^5,37^, correlates with gp27, while the C-terminal region is analogous to gp5 (**Sup.Fig.6g**). The upper part of the central spike adopts a β-barrel conformation, featuring the folding seen in the tube proteins, allowing the transition between the 6-fold symmetry of the tube initiator hexamer and the 3-fold symmetry of the spike trimer. The lower part of the central spike folds in a cone-shaped manner, creating a rigid structure stabilized by several integrated β-strands, and binds to the spike tip protein Pvc10, a homolog of gp5.4 in phage T4 (**Fig.2d**). We attempted to de-symmetrize the spike tip density but this was not possible, and no atomic model was built for Pvc10. Thus, the AlphaFold prediction of Pvc10 was docked into the tip density, following the β-strand folding of Pvc8 and using the structure of T4 gp5 and gp5.4 in their C-terminal regions as a reference. Some phages have been reported to puncture cell membranes with ion-loaded spikes^37,38^. To determine possible loading of ions in the spike tip of *Pl*PVC1, the sequence and predicted structure of Pvc10 were analyzed and compared to the sequence and structure of gp5.4. No conserved residues with potential involvement in iron coordination were found in Pvc10, in contrast to gp5.4, where an iron atom is coordinated by several histidine residues. However, two conserved residues in Pvc10, S34 and D49, correlate with two conserved residues in gp5.4, T25 and D41, which are believed to be involved in sodium binding (**Sup.Fig.6h**).

### *Pl*PVC1 plug

A helical density was identified within the cavity of the central spike, exposed toward the lumen of the inner tube (**Fig.2d**). Local refinement applying 3-fold symmetry allowed the structural identification of three copies of the plug protein Pvc6. Homologs of this protein could also be identified in corresponding regions of other particles^19,20,27^ (**Sup. Fig.7**, **Sup.Tabs.4-5**). A partial atomic model of Pvc6 could be constructed, from residues 24 to 51, a region that adopts an α-helical structure. Pvc6 features a trimeric hydrophobic inner core and a hydrophilic surface, which allows interaction with Pvc8 in its gp27-like region (**Sup.Fig.6i-6j**). Previous studies on plug homologs indicated that they are crucial for particle assembly and functionality^20,27^. To further validate this, a *Pl*PVC1ΔPvc6 mutant was generated and analyzed. No assembled particles could be purified (**Sup.Fig.6k**).

### *Pl*PVC1 trunk: inner tube

The tube of the *Pl*PVC1 particle is composed of three different tube proteins (Pvc7, Pvc5, Pvc1), which assemble a structure with inner and outer diameters of 45 Å and 80 Å, respectively (**Fig.1b**). The first layer of the tube, that contacts the central spike Pvc8, is formed by the tube initiator Pvc7, followed by a ring of Pvc5, and then continued by consecutive stacked layers of Pvc1 until the apical end (**Fig.1b**, **Fig.3a**).

**Figure 3.**
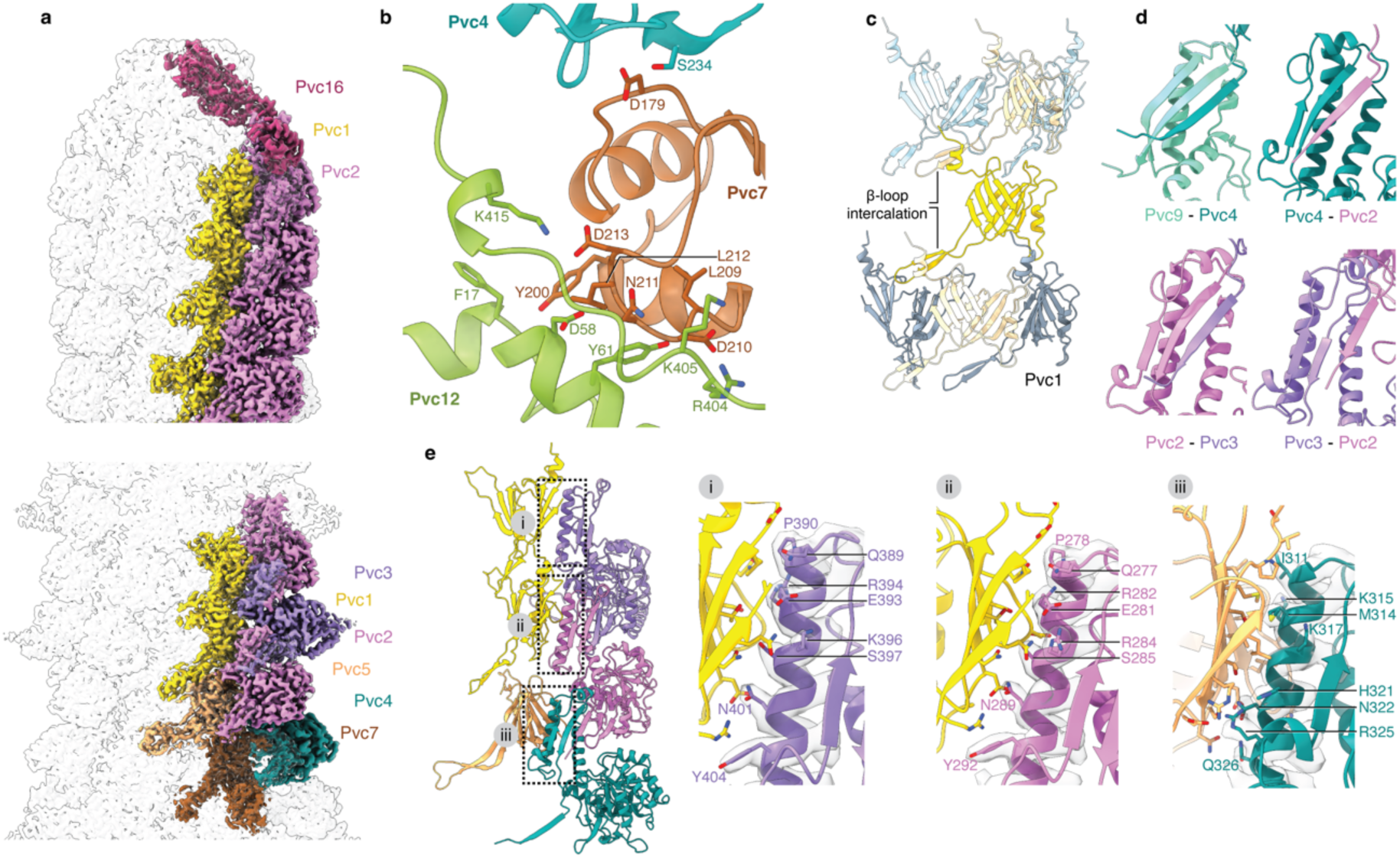
Organization and atomic model of the *Pl*PVC1 tube and sheath in its extended state. **(a)** Cryo-EM map of the extended *Pl*PVC1 particle with colored tube and sheath subunits, at cap level (*top*) and at baseplate level (*bottom*). **(b)** Zoom-in view of the main interactions of the LysM-like domain in Pvc7 with the sheath initiator Pvc4 and the baseplate wedge Pvc12. Residues involved in the main interactions are labeled and shown as sticks. **(c)** β-loop intercalations within Pvc1 subunits in different stacked layers of the tube. One central Pvc1 subunit is represented in yellow, interacting with two Pvc1 subunits in the layer above (light blue) and two in the layer below (grey). **(d)** Zoom-in view of the β-intercalation handshakes between sheath layers. The β-strand exchange is conserved along all layers, from baseplate to cap. **(e)** Conserved interactions between tube and sheath subunits, showing cryo-EM density and ribbon diagrams. Residues in the attachment helices of the different sheath subunits are labeled and shown as sticks. (i) Pvc3-Pvc1, (ii) Pvc2-Pvc1, and (iii) Pvc4-Pvc5.

Pvc7, Pvc5, and Pvc1 proteins share a common fold, similar to their corresponding homologs gp48, gp54, and gp19 in phage T4 (**Sup.Fig.8a-8b**). Additionally, Pvc7 features a LysM-like domain^39^, as gp53 in phage T4 and glue in pyocin R2. This domain is also present in the tube initiators in AFP, PVCpnf, and AlgoCIS (**Sup.Fig.8c**). The LysM-like domain extends in the C-terminal region of Pvc7 and interacts extensively with the baseplate wedge protein Pvc12 and the sheath protein Pvc4, playing an important role in the stabilization of the baseplate-trunk interface (**Fig.3b**).

The stacking of tube proteins relies on β-loop intercalations between them. Each Pvc1 subunit interacts with two subunits in the layer above and two in the layer below (**Fig.3c**). Following the same pattern, Pvc5 and Pvc7 subunits interact with Pvc1-Pvc7 and with Pvc5-Pvc8, respectively. This β-barrel-like arrangement lengthens the whole tube from central spike to cap, providing a compact and rigid structure. Notably, the inner surface of the tube lumen is negatively charged (**Sup.Fig.8d**), which in eCISs is believed to be involved in the efficient packing and release of cargoes loaded inside the trunk^19,27,40^.

### *Pl*PVC1 trunk: sheath

The sheath of the *Pl*PVC1 particle is formed by three different proteins (Pvc4, Pvc2, Pvc3) which surround the inner tube, expanding the outer diameter of the trunk to 162 Å (**Fig.1b**, **Fig.3a**). Pvc2, Pvc3, and Pvc4 share a common fold, similar to gp18 in phage T4 and sheath proteins in other eCIS (**Sup.Fig.9a-9b**). When compared to the other sheath proteins, Pvc3 presents an extra knob, formed by residues 64-117 and 225-278, which is believed to act as a fiber docking domain for the retracted fibers in the extended state of the particle (**Fig.1c**, **Sup.Fig.9a**).

The sheath is initiated by the sheath initiator Pvc4, which interacts with the sheath adaptor Pvc9, the LysM-like domain in Pvc7, Pvc5 in the first layer of the tube, and Pvc2 in the first layer of the sheath (**Sup.Fig.9c**). In the following levels, the sheath comprises alternate layers of Pvc2 and Pvc3, finishing with stacked layers of Pvc2 in the apical end, all assembled with a helical rise of 39.8 Å and a twist of 20.1° (**Fig.1b**, **Fig.3a**). The alternating Pvc2-Pvc3 pattern seems to be influenced by the assembly of the retracted fibers (**Fig.1b-1c**), as Pvc3 is the only sheath protein with a presumed fiber docking domain (**Sup.Fig.9a**). The exact layer at which Pvc3 terminates could not be determined due to length heterogeneity in a mixed population of particles.

The sheath assembly relies on β-strand intercalations, or handshakes, between sheath proteins. The first intercalation happens between the sheath adaptor Pvc9 and the sheath initiator Pvc4, which allows for the docking of the sheath and baseplate together (**Fig.3d**, **Sup.Fig.9c**). Analogously, Pvc4 interacts with Pvc2, which sequentially interacts with Pvc3. These consecutive handshakes propagate in each sheath layer, from baseplate to cap, stabilizing the sheath assembly in the *Pl*PVC1 particle (**Fig.3d**).

The sheath and tube proteins interact along the particle via generally conserved interactions^19,27,40^. Pvc2, Pvc3, and Pvc4 feature an attachment helix through which interactions with the tube proteins Pvc1 and Pvc5 occur (**Fig.3e**). The tube-sheath interplay seems to be driven by the specific distribution of positive and negative charges at the contacting interfaces, contributing to the stabilization of the particle in its extended state.

### *Pl*PVC1 terminal cap

The extended *Pl*PVC1 particle terminates with the cap complex at the apical end. This complex is composed of six monomers of the protein Pvc16 (**Fig.1c**, **Fig.4a**). Each Pvc16 monomer consists of two main domains (N-terminal and C-terminal) connected by a middle loop with a β-strand (**Fig.4b**). Pvc16 N-terminal domain has a similar fold to gp15 from phage T4^41^ but contains an extra α-helix that allows the closure of the inner tube down to a diameter of 6.7 Å, as has also been observed for Pvc16 of PVCpnf^27^ and Afp16^19^ (**Fig.4c**, **Sup.Fig.10a-10b**). There are multiple interactions between the Pvc16 subunits and tube and sheath proteins. The N-terminal domain of Pvc16 interacts with two different subjacent Pvc1 neighboring subunits in their N-terminal region, leading to a unique conformation of Pvc1 in the apical layer (**Fig.4d**, **Sup.Fig.10c**). The middle loop of each Pvc16 monomer interacts via β-strand intercalation with the C-terminal region of a Pvc2 subunit in the top layer of the sheath, while the C-terminal domain of Pvc16 makes a turn and interacts with the N-terminal region of the adjacent Pvc2 subunit (**Fig.4e**, **Sup.Fig.10c**). These interactions allow for the docking of Pvc16 into the upmost sheath layer in a handshake manner, closing and stabilizing the particle in its extended state.

**Figure 4.**
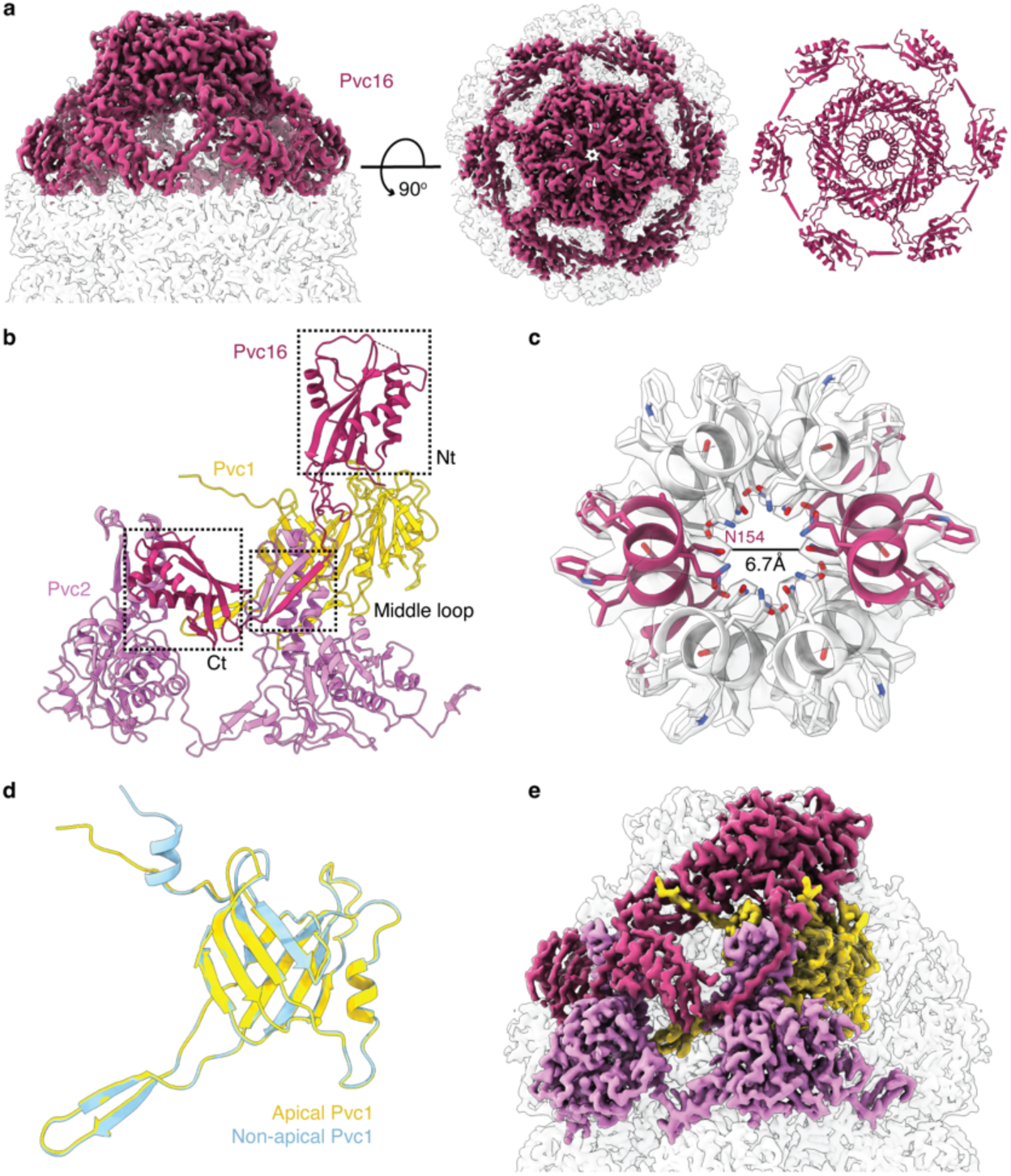
Organization and atomic model of the *Pl*PVC1 cap in its extended state. **(a)** Side and top views of the cryo-EM map of the *Pl*PVC1 terminal cap, with colored Pvc16 subunits, together with top view of the atomic model of the Pvc16 complex in ribbon diagram. **(b)** Zoom-in view of a Pvc16 monomer interacting with Pvc1 and Pvc2 in the upmost layer of the particle in its extended state. The N-terminal domain (Nt), middle β-strand loop, and C-terminal domain (Ct) of Pvc16 are marked with dashed rectangles. **(c)** Central ring of the terminal cap complex, showing cryo-EM density and ribbon diagrams, with fitted α-helices (residues 149-159 shown as sticks). Two opposite α-helices are colored in dark magenta as reference. The amino acid N154 was used to measure the 6.7 Å aperture of the apical part of the *Pl*PVC1 particle. **(d)** Conformational comparison of the apical Pvc1 tube subunit in the top layer (yellow) with the non-apical Pvc1 tube subunit in the rest of the layers (blue). **(e)** Side view of the cryo-EM map of the *Pl*PVC1 terminal cap, showing the conserved β-intercalation handshake between Pvc16 and Pvc2 subunits, together with the interaction between Pvc16 and the apical Pvc1 subunit.

### *Pl*PVC1 contracted sheath and baseplate of the contracted particle

In order to investigate the conformational changes that occur when particles get activated and sheath contraction is initiated, the contraction process was mimicked *in vitro* by exposing purified particles to 3 M urea^31,32^ (**Fig.1d**, **Sup.Fig.2c**). As in other CISs^5,11,19,27,40^, the sheath undergoes conformational changes after contraction, without losing the handshakes between subunits, which seemingly maintains the integrity of the sheath (**Fig.5a-5b**, **Sup.Fig.11a**). Contraction leads to vertical compression of the sheath, with a helical rise of 17.5 Å and a twist of 31.7°, and to an expansion in both the outer and inner diameters, from 162 Å to 220 Å in the former, and from 80 Å to 110 Å in the latter (**Fig.5b**, **Sup.Fig.11b**). This is driven by conformational transitions in the sheath proteins, which rearrange their C- and N-termini by rigid-body rotation, compared to their conformation in extended state (**Sup.Fig.11c**). These rearrangements allow each sheath monomer to slide on top of the adjacent one, opening the diameter of the particle and enabling tube-sheath detachment and tube ejection for target cell membrane perforation.

**Figure 5.**
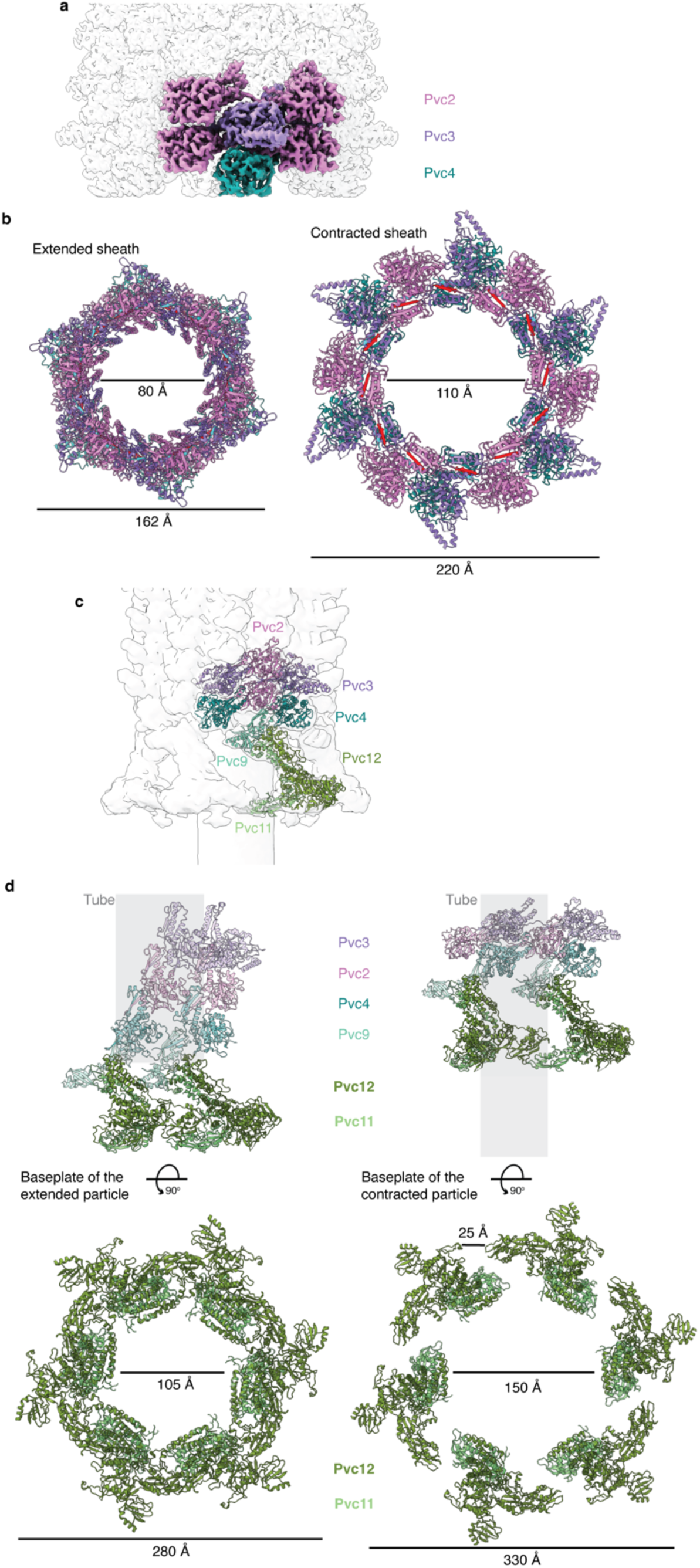
Organization and atomic model of the *Pl*PVC1 sheath and baseplate in its contracted state. **(a)** Cryo-EM map of the *Pl*PVC1 sheath in contracted state, with colored sheath subunits. **(b)** Top view of the ribbon diagrams of sheath subunits Pvc2, Pvc3, and Pvc4 in extended (*left*) and contracted (*right*) states, with diameter measurements. The β-strands involved in subunit intercalation are colored in red (β-strands in C-terminal region) and cyan (β-strands in N-terminal region). **(c)** Cryo-EM map of the *Pl*PVC1 baseplate in its contracted state, filtered to 10 Å, with fitted Pvc2, Pvc3, Pvc4, Pvc9, Pvc11, and Pvc12 subunits. Map-model Correlation Coefficient = 0.65. **(d)** *Top*: Side view of the ribbon diagrams of two adjacent baseplate wedges (Pvc11 and Pvc12), together with first layers of the sheath (Pvc9, Pvc4, Pvc2, and Pvc3), in extended (*left*) and contracted states (*right*). Sheath proteins are represented in transparent colors. Baseplate wedges are represented in solid colors. Tube is represented in light grey. *Bottom*: top view of the ribbon diagrams of baseplate wedges (Pvc11 and Pvc12) in extended (*left*) and contracted (*right*) states, with measurements for diameter and separation of the wedges.

The density map of the baseplate of the contracted *Pl*PVC1 particle could be solved at a resolution ranging from 4 Å to 10 Å, corresponding to areas closer to the sheath and periphery of the wedges, respectively (**Sup.Fig.3-4**, **Sup.Tab.3**). An atomic model could not be built *de novo* for the baseplate of the contracted particle, but a partial atomic model was obtained by rigid-body fitting of the sheath adaptor Pvc9 and baseplate wedges Pvc11 and Pvc12 into the density (**Fig.5c**). This fitted model suggested a rearrangement of Pvc9, Pvc11, and Pvc12 in the baseplate of the contracted particle, compared to its conformation in the extended particle, leading to the opening of the diameter of the wedges (**Fig.5d**). The expansion and lateral dissociation of the baseplate wedges after contraction has also been reported in the AFP particle^19^ and pyocin R2^11^. As shown for gp25 in phage T4^5^, the sheath adaptor Pvc9 is believed to play an important role in particle contraction by transducing the contraction signal from the baseplate to the sheath, launching the subsequent conformational changes in the sheath proteins.

### *Pl*PVC1 fiber

*Pl*PVC1 presents a set of six tail fibers, arranged in a retracted manner around the extended particle (**Fig.1b**). Each fiber is composed of three intertwined copies of the Pvc13 protein and exhibits an arched topology, in which the C-terminus is folded toward the middle of the fiber (**Sup.Fig.12a-12b**). Pvc13 sequence analysis showed an organization of the fiber into three main parts: the N-terminal region containing helical motifs with homology to fibers in other CISs^42^, the central region with repetitive motifs homologous to adenoviruses fibers, and the C-terminal region with homology to host-binding domains of short tail fibers from bacteriophages (**Sup.Fig.12c**).

The AlphaFold model of the fiber, a trimer of Pvc13, was fitted into the density determined for the fiber in retracted conformation. The resolution was sufficient for recognizing domain segments of the fiber and for confident fitting of the AlphaFold model into the density (**Fig.6a**, **Sup.Fig.12b**). Local refinement over the area of interaction between baseplate and fiber allowed for atomic modeling of this region (residues 29-55 in Pvc13 and residues 622-641, 686-708, 882-956 in Pvc12) (**Fig.6b**). The interaction between baseplate and fiber happens between the C-terminal region of Pvc12, in the periphery of the baseplate wedge, and the N-terminal region of the Pvc13 trimer, which folds into two conserved α-helices (**Fig.6b**, **Sup.Fig.12d**). This interaction likely contributes to the orientation of the fiber in a retracted conformation in the extended state of the particle.

**Figure 6.**
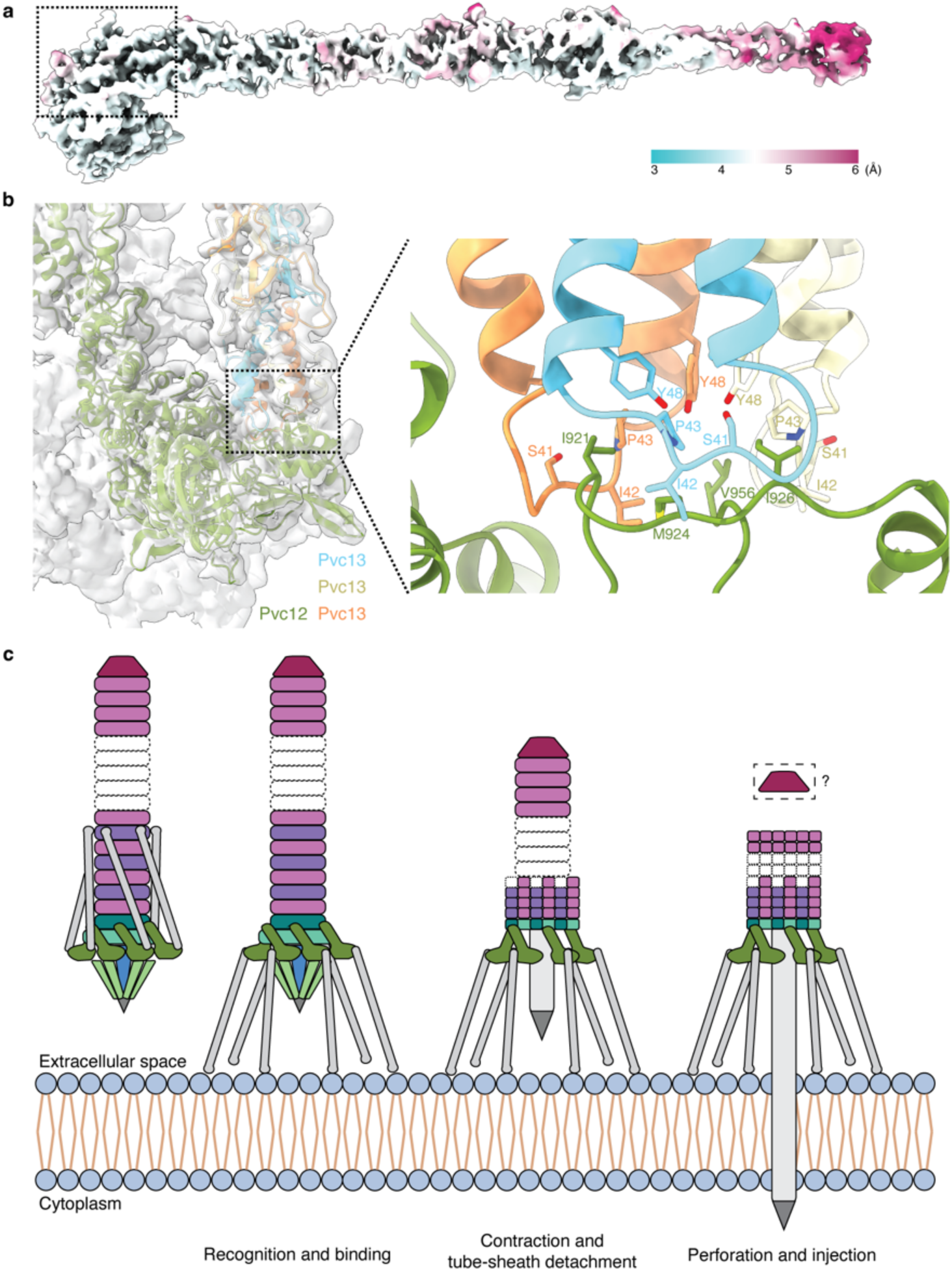
Structure of the *Pl*PVC1 fiber and schematic model for target recognition and contraction. **(a)** Cryo-EM map of the *Pl*PVC1 fiber colored by local resolution. The dashed rectangle marks the region where local refinement was applied to refine the interaction between baseplate and fiber. **(b)** Cryo-EM map and atomic model of the interaction between baseplate and fiber. Pvc12 is colored in dark green. Each chain of Pvc13 is colored in cyan, yellow, and orange, respectively. Residues involved in the baseplate-fiber interaction are labeled and shown as sticks. **(c)** Schematic representation of the proposed mechanism for target cell recognition, particle binding, particle contraction, and perforation of the target cell membrane. The particle recognizes the target cell via the tail fiber network, leading to particle attachment and contraction triggering, which terminates with the translocation of the spike through the target cell membrane.

## DISCUSSION

Bacteria have evolved specialized contractile injection systems to invade and modulate target cells, all evolutionary related to bacteriophages tails^4^. To date, several CISs have been broadly studied, and their high-resolution structures have been described^5,7,8,11,19,20,27,40,42^. The work here presents the single-particle cryo-EM structure of a PVC particle from *Photorhabdus luminescens DJC* (*Pl*PVC1), in both its extended and contracted states, highlighting its evolutionary relationship with other CISs, and provides a molecular framework for understanding its mechanism of action.

The *Pl*PVC1 particle has a contractile trunk with conserved handshakes and β-loop intercalations between the stacked sheath and tube subunits and is stabilized at the apical end by the terminator cap complex. The hexagonal baseplate in *Pl*PVC1 is assembled in a 1:1:1 stoichiometry around the central spike, which is sharpened by the spike tip. Interestingly, the *Pl*PVC1 particle features a longer sheath adaptor which protrudes on top of the baseplate wedges. Although analysis of the extra domain in the sheath adaptor was performed, the functionality of this protrusion remains ambiguous and requires further investigation. Two other remarkable features of the *Pl*PVC1 particle are the presence of a plug protein in the cavity of the central spike exposed toward the lumen of the inner tube, and the existence of baseplate extensions forming a cage around the central spike. The functionality of the cage remains unknown, but our study supports Xu *et al*.’s hypothesis^20^ on possible contribution of the cage to particle-cell attachment. In addition, our results also reinforce the importance of the plug protein in particle assembly, as no assembled particles were observed in the *Pl*PVC1ΔPvc6 mutant sample.

The structure of a PVC fiber has not been previously described at high resolution. In this study, the density map of the *Pl*PVC1 fiber in retracted conformation is solved at a resolution range of 4 Å to 6 Å, the AlphaFold model of the fiber is confidently fitted into the density, and the atomic interactions between baseplate and fiber are modeled. Sequence analysis of the *Pl*PVC1 fiber showed divergence from phage tail fibers in the presence of homologous regions with fibers from eukaryotic viruses (adenoviruses), which suggests that the fiber protein Pvc13 could be a fusion protein derived from phage tail fibers and adenovirus fibers and reinforces the hypothesis of PVC particles targeting eukaryotic organisms^27,28^.

Extended and contracted structures of other eCISs have been reported, providing insight into the mechanism of contraction^11,19,20,27,40^. However, high-resolution structures of the baseplates in the contracted state remain incomplete in most cases. Here, we present a cryo-EM structure of the baseplate of the contracted *Pl*PVC1 particle, facilitating our understanding of the contraction mechanism in this particle and allowing the comparison with other eCISs. *In vitro* contraction of the *Pl*PVC1 particle established the molecular framework for studying protein rearrangements in the contracted state and for comparing it with the metastable extended state.

In a similar fashion to phage T4^5,32,43–45^, the contraction signal is believed to be sensed at the level of the fibers, after recognition of specific receptors on the target cell membrane^28,46^. Subsequently, changes in the orientation of the fibers are transmitted to the baseplate through Pvc12, leading to expansion of the wedges and pivoting toward the periphery of the particle. Afterwards, contraction propagates toward the sheath, until reaching the terminal part of the particle. The sheath adaptor Pvc9 plays a crucial role as a connector between the dilated baseplate and the sheath, as also seen in phage T4^5^, AFP particle^19^, and pyocin R2^11^. Rearrangements in the sheath adaptor trigger further conformational changes along the sheath, without affecting its structural integrity. The widening of the contracted sheath leads to sheath compaction, tube-sheath detachment, and tube ejection, with final perforation of the target cell membrane to inject the payload into the target cell (**Fig.6c**). The puncturing end of the tube, which is sharpened by the central spike and spike tip, is crucial for the perforation. Both the rigidity of the spike and the conical shape of the tip play important roles in the piercing process. Indeed, the rigid spike is believed to translocate through the membrane without major unfolding and, in some phages, loaded with ions^37,38^. Our analysis of the sequence and predicted structure of the spike tip protein Pvc10 suggests the hypothesis of potential loading of sodium ions in the tip, which may contribute to target cell membrane digestion during piercing.

Upon perforation, PVCs can translocate toxins through eukaryotic cell membranes^21,22^. Functional studies employing PVCs have demonstrated specific delivery of protein cargoes into selected target cells, together with efficient reprograming of receptor recognition by tail fibers^28,29^. The ability of modified PVCs to recognize and deliver payloads to specific targets could be harnessed for the development of novel targeted drug delivery systems. The identification of key structural features involved in contraction and target selection is crucial for broadening the opportunities to engineer PVCs with modified host recognition properties and customized cargo-loading capabilities. Future research should focus on elucidating the full range of payloads delivered by these systems and optimizing PVCs as biotechnological tools for biomedical use.

In conclusion, by elucidating structural features of the novel *Pl*PVC1 particle, together with transitions between extended and contracted states, this study enhances our understanding of contractile injection systems and their evolutionary links to bacteriophages. The findings presented here deepen our knowledge of bacterial nanomachines and lay more groundwork for harnessing PVCs in biomedical and agricultural applications. Further studies investigating their functional mechanisms and potential engineering approaches will be crucial for unlocking their full potential.

## METHODS

### Experimental model and subject details

*E. coli* strains were cultured aerobically in LB medium [1% (w/v) NaCl; 1% (w/v) tryptone; 0.5% (w/v) yeast extract] at 37 °C. *E. coli* HST08 strain (Stellar chemically competent cells) was used for DNA manipulation, and *E. coli* BL21Star(DE3) was used for protein expression.

*Photorhabdus luminescens DJC* strain (TT01-RifR) was obtained from the lab of Prof. Dr. Ralf Heermann (Johannes Gutenberg University of Mainz, Germany). This strain was cultivated aerobically in CASO medium [5% (w/v) NaCl; 1.5% (w/v) peptone from casein; 0.5% (w/v) peptone from soymeal] at 30 °C.

For preparation of agar plates, 1% (w/v) agar was added to the respective medium. Antibiotics were used as follows: ampicillin 100 μg/mL; chloramphenicol 34 μg/mL; rifampicin 50 μg/mL.

### Cloning of *Pl*PVC1 encoding operon

The *pvc* operon 1 from *Photorhabdus luminescens DJC* (*PluDJC_08925* to *PluDJC_08830*) was amplified by PCR from genomic DNA and cloned into pBAD33 plasmid (arabinose-inducible promoter, chloramphenicol resistance), previously linearized by PCR, using primers with an overlap with the first and last ORF in the *Pl-pvc1* cluster. After DNA fragment purification, insert and vector were mixed in a 1:1 ratio and incubated with In-Fusion® Snap Assembly Master Mix (Takara) for 15 minutes at 50 °C. *E. coli* Stellar competent cells were transformed with 2.5 μL of In-Fusion reaction and incubated overnight. Positive clones were screened by colony PCR and restriction enzyme digestion (BamHI and BsaI) after plasmid extraction. The full plasmid sequence was verified by Next Generation Sequencing. PCR reactions were performed with Platinum^TM^ SuperFi^TM^ PCR Master Mix (Invitrogen), and DNA fragment purification was carried out using QIAGEX II Gel Extraction kit (Qiagen).

### *Pl*PVC1 particle expression

Verified pBAD33-PluDJC_08925-08830 plasmid was transformed into *E. coli* BL21Star(DE3) electrocompetent cells. After selection, cells were grown overnight at 37 °C in 10 mL LB medium supplemented with chloramphenicol. The following day, 1 L of LB medium supplemented with chloramphenicol, was inoculated with overnight culture, and protein expression was induced with 0.2% L-Arabinose at OD_600_ 0.7. Cells were incubated for 24 hours at 18 °C with slow agitation (80 rpm). Cells were harvested at 5,000 rpm for 20 minutes at 4 °C. The cell pellet was resuspended in 50 mL of cold milli-Q water. Washing was carried out by centrifugation at 4,000 rpm for 15 minutes at 4 °C. The final pellet was flash-frozen in liquid nitrogen for 5 minutes and stored at −20 °C.

### *Pl*PVC1 particle purification

Bacterial cell pellets were lysed in 25 mL of lysis buffer^27^ (25 mM Tris pH 7.4, 140 mM NaCl, 3 mM KCl, 200 μg/mL lysozyme, 50 μg/mL DNase I, 0.5% Triton X-100, 5 mM MgCl_2_, 1x protease inhibitor) for 1 hour at 37 °C. The cell lysate was cleared by two rounds of centrifugation (6,000*g* for 30 minutes at 4 °C, and 30,000*g* for 30 minutes at 4 °C). The particles were pelleted by ultracentrifugation at 100,000*g* for 1 hour at 4 °C. The particle pellet was resuspended overnight in 2 mL of Tris-salt buffer^20^ (20 mM Tris pH 7.5, 150 mM NaCl). The resuspension was applied on an iodixanol-based gradient (10%-40%) and subjected to ultracentrifugation at 100,000*g* for 20 hours at 4 °C. The gradient was divided into 12 fractions and each fraction was checked for presence of *Pl*PVC1 particles by negative-staining electron microscopy. The fractions containing the particles were buffer-exchanged (from iodixanol to Tris-salt buffer) via dialysis in 20 kDa MWCO cassettes for 6 days at 4 °C. After dialysis, particles were pelleted by ultracentrifugation at 100,000*g* for 1 hour at 4 °C. The pellet was finally resuspended in 100 μL of Tris-salt buffer and cleared by centrifugation 10,000*g* for 5 min at 4 °C. The supernatant containing the *Pl*PVC1 particles was stored at 4 °C for short-term use.

### Mass spectrometry sample preparation

100 µL of room-temperature 50 mM ammonium bicarbonate was added to 20 µg (in ∼5 µL) of purified *Pl*PVC1 sample. Following this, 0.5 µg of sequencing-grade trypsin was added, and the sample was incubated overnight at 25 °C with gentle mixing. The digest was reduced and alkylated by concomitant addition of tris(2-carboxyethyl)phosphine and chloroacetamide to final concentrations of 10 mM, and incubating at 30 °C for 30 min. The sample was clarified through a 0.45 µm spin filter, and peptides were purified via high-pH C18 StageTip procedure. To this end, C18 StageTips were prepared in-house, by layering four plugs of C18 material (Sigma-Aldrich, Empore SPE Disks, C18, 47 mm) per StageTip. Activation of StageTips was performed with 100 μL 100% methanol, followed by equilibration using 100 μL 80% acetonitrile (ACN) in 200 mM ammonium hydroxide, and two washes with 100 μL 50 mM ammonium hydroxide. The sample was basified to pH >10 by addition of one tenth volume of 200 mM ammonium hydroxide, and loaded on two StageTips. Subsequently, StageTips were washed twice using 100 μL 50 mM ammonium hydroxide, after which peptides were eluted using 80 µL 25% ACN in 50 mM ammonium hydroxide. The samples were dried to completion using a SpeedVac at 60 °C. Dried peptides were dissolved in 20 μL 0.1% formic acid (FA) and stored at −20 °C until analysis using mass spectrometry.

### Mass spectrometry data acquisition

Around 2 µg of digested proteins (∼500 ng of peptide) were analyzed per injection, with three technical replicates. All analyses were performed on an EASY-nLC 1200 system (Thermo) coupled to an Orbitrap Exploris 480 mass spectrometer (Thermo). Samples were analyzed on 20 cm long analytical columns, with an internal diameter of 75 μm, and packed in-house using ReproSil-Pur 120 C18-AQ 1.9 µm beads (Dr. Maisch). The analytical column was heated to 40 °C, and elution of peptides from the column was achieved by application of gradients with stationary phase Buffer A (0.1% FA) and increasing amounts of mobile phase Buffer B (80% ACN in 0.1% FA). The primary analytical gradients ranged from 5 %B to 38 %B over 60 min, followed by a further increase to 48 %B over 5 min to elute any remaining peptides, and finally a washing block of 15 min. Ionization was achieved using a NanoSpray Flex NG ion source (Thermo), with spray voltage set at 2 kV, ion transfer tube temperature to 275 °C, and RF funnel level to 40%. Full scan range was set to 300-1,300 m/z, MS1 resolution to 120,000, MS1 AGC target to “200” (2,000,000 charges), and MS1 maximum injection time to “Auto”. Precursors with charges 2-6 were selected for fragmentation using an isolation width of 1.3 m/z and fragmented using higher-energy collision disassociation (HCD) with normalized collision energy of 25. Monoisotopic Precursor Selection (MIPS) was enabled in “Peptide” mode. Precursors were prevented from being repeatedly sequenced by setting dynamic exclusion duration to 80 s, with an exclusion mass tolerance of 15 ppm and exclusion of isotopes. MS/MS resolution was set to 30,000, MS/MS AGC target to “200” (200,000 charges), MS/MS intensity threshold to 360,000 charges/second, MS/MS maximum injection time to “Auto”, and number of dependent scans (TopN) to 13.

### Mass spectrometry data analysis

All RAW files were analyzed using MaxQuant software (v1.5.3.30). Default MaxQuant settings were used, with exceptions outlined below. For generation of the theoretical spectral library, all expected full-length PVC protein sequences were entered into a FASTA database. Digestion was performed using “Trypsin/P” (default), allowing up to 3 missed cleavages. Minimum peptide length was set to 6, and maximum peptide mass to 6,000 Da. Protein N-terminal acetylation (default), oxidation of methionine (default), deamidation of asparagine and glutamine, and peptide N-terminal glutamine to pyroglutamate, were included as potential variable modifications, with a maximum allowance of 3 variable modifications per peptide. Modified peptides were stringently filtered by setting a minimum score of 100 and a minimum delta score of 50. First search mass tolerance was set to 10 ppm, and maximum charge state of considered precursors to 6. Label-free quantification (LFQ) was enabled, with “Fast LFQ” disabled. Second peptide search was disabled. Matching between runs was enabled with a match time window of 1 min and an alignment time window of 20 min. Data was filtered by posterior error probability to achieve a false discovery rate of <1% (default), at the peptide-spectrum match, protein assignment, and site-decoy levels.

### *Pl*PVC1 particle contraction

Particle contraction was performed via dialysis in 3 M urea^31,32^. 70 μL of purified *Pl*PVC1 sample were placed in a mini dialysis cassette of 20 kDa MWCO. The sample was first dialyzed for 4 hours in 3 M urea, pH 7.5, at 4 °C, and then dialyzed in Tris-salt buffer for another 4 hours at 4 °C. The contracted sample was stored at 4 °C until further use.

### Electron microscopy

For negative-staining electron microscopy, 4 μL of *Pl*PVC1 samples were applied onto glow-discharged (30 sec, 15 mA, in a Leica ACE 200) copper grids coated with a continuous carbon layer, then washed 3 times with 50 μL milli-Q water, and finally stained with 2% uranyl acetate. The grids were dried at room temperature and imaged on a Morgagni 268 transmission electron microscope operated at 100 kV.

### Cryo-EM grids preparation

For cryo-EM, 3 μL of *Pl*PVC1 samples were applied to glow-discharged (10 sec, 5 mA, in a Leica ACE 200) Quantifoil grids (R2/2, 200 mesh Gold, coated with a 2 nm continuous carbon layer), and plunge-frozen into liquid ethane pre-cooled with liquid nitrogen, using a Vitrobot Mark IV (FEI, Thermo Fisher Scientific) at 4 °C and 100% humidity.

### Cryo-EM data collection, image processing, and refinement

The cryo-EM grids were screened on a Glacios cryo-TEM at 200 kV (Thermo Fisher Scientific), equipped with a Falcon 3 Direct Electron Detector. Data acquisition was performed on a Titan Krios G2 at 300 kV (Thermo Fisher Scientific), paired with a Falcon 4i Direct Electron Detector and SelectrisX energy filter.

Micrographs were collected using the semi-automated acquisition program EPU (FEI, Thermo Fisher Scientific) at 105,000x magnification, with a calibrated pixel size of 1.2 Å and a defocus range of −0.6 to −2.0 μm.

All datasets were processed using cryoSPARC^47^ v4.3.0 to v4.6.2. First, patch motion correction was used to estimate and correct for full-frame motion and sample deformation (local motion). Patch contrast transfer function (CTF) estimation was used to fit local CTF to micrographs. Micrographs were manually curated to remove low-quality data based on ice thickness, local-motion distances, and CTF-fit parameters. Particles were picked using Topaz particle picking^48^. First, Topaz was trained with a manually picked set of particles. Then, Topaz Extract was used with the pre-trained model and a pre-tested particle threshold value. This procedure was performed equally for the baseplate of the extended particle, the cap, and the baseplate of the contracted particle.

#### Baseplate

After Topaz particle picking and picks inspection, particles were extracted with a box size of 700 pixels and Fourier-cropped to 352 pixels. One round of 2D classification was performed followed by *ab initio* 3D reconstruction. The 3D density was refined by non-uniform refinement, with imposed C6 symmetry. After particle re-extraction with full box size (700 pixels), non-uniform refinement, with imposed C6 symmetry, was applied with a dynamic mask to obtain a high-resolution map.

#### Central spike

The C6-symmetrized 3D volume of the baseplate was shifted toward the central spike region, and particles were re-extracted with a box size of 360 pixels. After 3D reconstruction and refinement, particles were subjected to symmetry expansion (total copies = 6). One round of 3D classification, with a focus mask around the central spike region, was performed. The density of the one class showing clear trimeric symmetry was refined by non-uniform refinement with C3 symmetry imposed. Duplicated particles were removed, and final high-resolution map of the central spike region was obtained by non-uniform refinement with imposed C3 symmetry.

#### Fiber

Particles from the binned C6-symmetrized 3D volume of the baseplate were re-extracted with a box size of 560 pixels and a binning factor of 1.25x. After 3D reconstruction and refinement, particles were subjected to symmetry expansion (total copies = 6). Two rounds of 3D classification, with focus mask around the fiber, were performed. Classes showing clear density in the masked area were refined by local refinement without symmetry imposition (C1), using the same mask applied during the 3D classifications.

#### Cap

After Topaz particle picking and picks inspection, particles were extracted with a box size of 560 pixels and Fourier-cropped to 288 pixels. One round of 2D classification was performed followed by *ab initio* 3D reconstruction. The 3D density was refined by non-uniform refinement, with imposed C6 symmetry. After particle re-extraction with full box size (560 pixels), non-uniform refinement with imposed C6 symmetry was applied with a dynamic mask to obtain a high-resolution map.

#### Baseplate of the contracted particle

After Topaz particle picking and picks inspection, particles were extracted with a box size of 700 pixels and Fourier-cropped to 352 pixels. One round of 2D classification was performed followed by *ab initio* 3D reconstruction. The 3D density was refined by heterogenous and non-uniform refinement, with imposed C6 symmetry. Particles were subjected to symmetry expansion (total copies = 6). One round of 3D classification, with focus mask around one baseplate wedge, was performed. The class showing clear density in the masked area was refined by local refinement without symmetry imposition (C1), using the same mask used for the 3D classification. Duplicated particles were removed, and the 3D density was refined by homogenous refinement with imposed C6 symmetry. After particle re-extraction with full box size (700 pixels), homogenous refinement with imposed C6 symmetry was applied to obtain a higher-resolution map.

#### Contracted sheath

The binned C6-symmetrized 3D volume of the baseplate of the contracted particle was subjected to local refinement, with imposed C6 symmetry and focus mask surrounding the first layers of the sheath immediately after the baseplate. One round of 3D classification, using the same focus mask, was performed. The class showing clear density in the masked area was refined by local refinement with imposed C6 symmetry. After particle re-extraction with full box size (700 pixels), local refinement with imposed C6 symmetry was applied to obtain a high-resolution map.

All the applied masks were created in UCSF ChimeraX v1.8^49^ and processed in cryoSPARC^47^. For all datasets, the number of micrographs, total exposure values, particles used for final refinement, map resolution, and other values during data processing are summarized in **Supplementary Table 3**. Cryo-EM data processing workflow and map resolutions with GSFSC curves are summarized in **Supplementary Figures 3-4**.

### Model building

The initial models of each protein in the *Pl*PVC1 particle were predicted using AlphaFold2^50^. Starmap^51^ v1.1.75 was used for automated building of the AlphaFold-predicted models in the density maps. Starmap results were inspected and manually adjusted in ISOLDE^52^ and Coot^53^. Atomic models were then refined against the corresponding maps using phenix.real_space_refine^54^ with secondary structure restraints and geometry restraints. Several iterations of phenix.real_space_refine, followed by manual adjustments in ISOLDE and Coot, were performed until convergence. Atomic models of Pvc6 and interaction between Pvc12 and Pvc13 were partially built due to density limitations. Atomic model of the baseplate of the contracted particle was generated by rigid-body fitting of the baseplate wedge proteins and the sheath adaptor into the solved density. A summary of the model refinement and validation statistics can be found in **Supplementary Table 3**.

### Bioinformatics analysis

Multiple sequence alignments (MSA) were performed using Clustal Omega^55^ and visualized using ESPript 3.0^56^. DALI web server^35^ and Foldseek Search Server^36^ were used for structural analysis and comparison. Protein interaction interfaces were predicted using the parameter-free geometric deep learning method PeSTo^33,34^ (Protein Structure Transformer). ConSurf Server^57^ was used for conservation analysis of sequence profiles.

## DATA AND SOFTWARE AVAILABILITY

The cryo-EM density maps and the corresponding atomic coordinates were deposited in the Electron Microscopy Data Bank (EMDB) and in the Protein Data Bank (PDB), respectively. The accession codes are listed as follows: baseplate (EMD-53137, 9QGL); central spike (EMD-53138, 9QGM); cap (EMD-53139, 9QGN); fiber and baseplate-fiber interaction (EMD-53140, 9QGO); contracted sheath (EMD-53141, 9QGP); baseplate of the contracted particle (EMD-53143). The mass spectrometry proteomics data have been deposited to the ProteomeXchange Consortium via the PRIDE^58^ partner repository with the dataset identifier PXD060336.

## SUPPLEMENTARY INFORMATION

Supplementary figures 1 to 12.

Supplementary tables 1 to 5.

## ACKNOWLEDGEMENTS

The Novo Nordisk Foundation Center for Protein Research is supported financially by the Novo Nordisk Foundation (NNF14CC0001). This work was supported by an NNF Hallas-Møller Emerging Investigator grant (NNF17OC0031006), an NNF Hallas-Møller Ascending Investigator grant (NNF23OC0081528), and an LF Ascending Investigator grant (R434-2023-289) to NMIT, who is a member of the Integrative Structural Biology Cluster (ISBUC) at the University of Copenhagen. This work was also supported by a fellowship from laCaixa Foundation (ID 100010434) with code LCF/BQ/EU21/11890147 to LMA. We acknowledge the Danish Cryo-EM Facility at the Core Facility for Integrated Microscopy (CFIM) at the University of Copenhagen for support during data collection. We acknowledge the Big Data Management Platform at Novo Nordisk Foundation Center for Protein Research for the computational resources.

## AUTHOR CONTRIBUTIONS

LMA, NMIT, and EMSR conceived the project. LMA performed cloning, particle expression, and particle purification and contraction. IAH performed mass spectrometry and analyzed the data, in consultation with MLN. LMA and ARE prepared cryo-EM grids and collected cryo-EM data, with assistance of TP and NS. LMA and ARE processed the cryo-EM data and determined the structures presented in this study. LMA performed the bioinformatic analysis. NMIT and LMA acquired the financial support for the project. LMA wrote the manuscript and prepared the figures, with input from all authors. All authors contributed to the revision of the manuscript.

