## SupplementaryInformation for "Structural characterization of an extracellular contractile injection system from *Photorhabdus luminescens* in extended and contracted states"

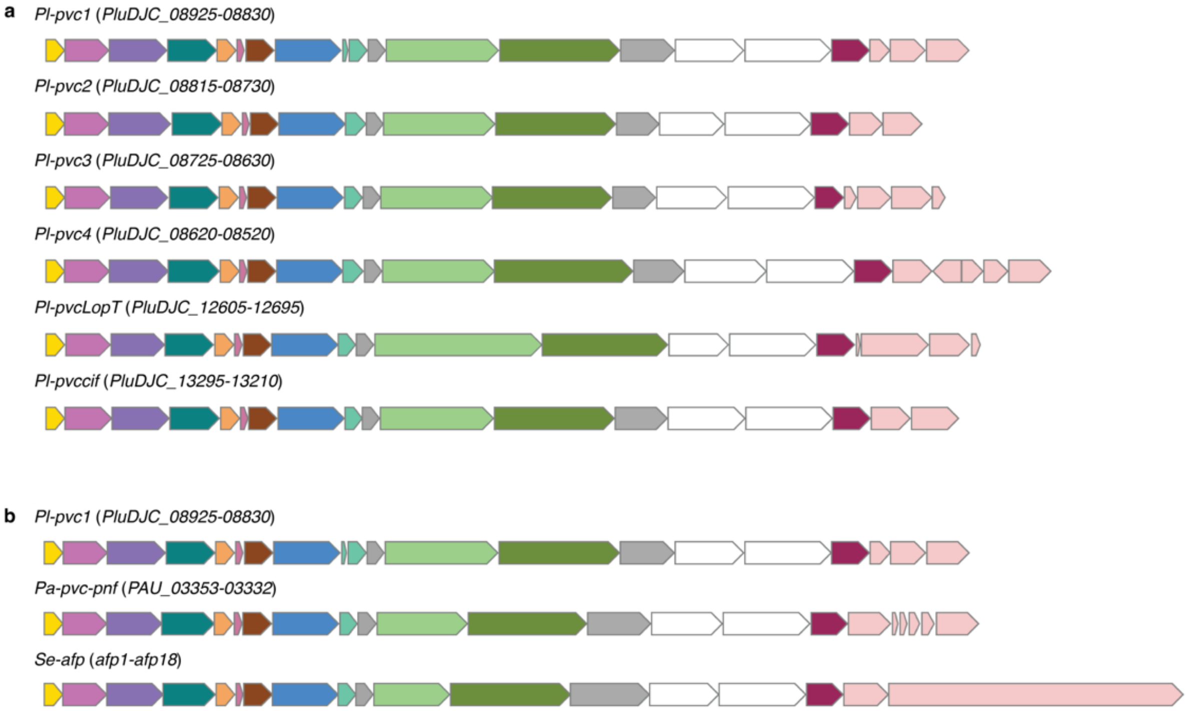


**Supplementary Figure 1. Comparison of *pvc* operons in *Photorhabdus luminescens DJC*, *pvc-pnf* operon, and *afp* gene cluster. (a)** Schematic representation of the six *pvc* operons in the genome of *P. luminescens DJC*. **(b)** Schematic representation of the *pvc* operon 1 from *P. luminescens DJC*, the *pvc-pnf* operon from *P. asymbiotica*, and the *afp* gene cluster from *S. entomophila*. The different genes are colored according to **Figure 1** in this manuscript.

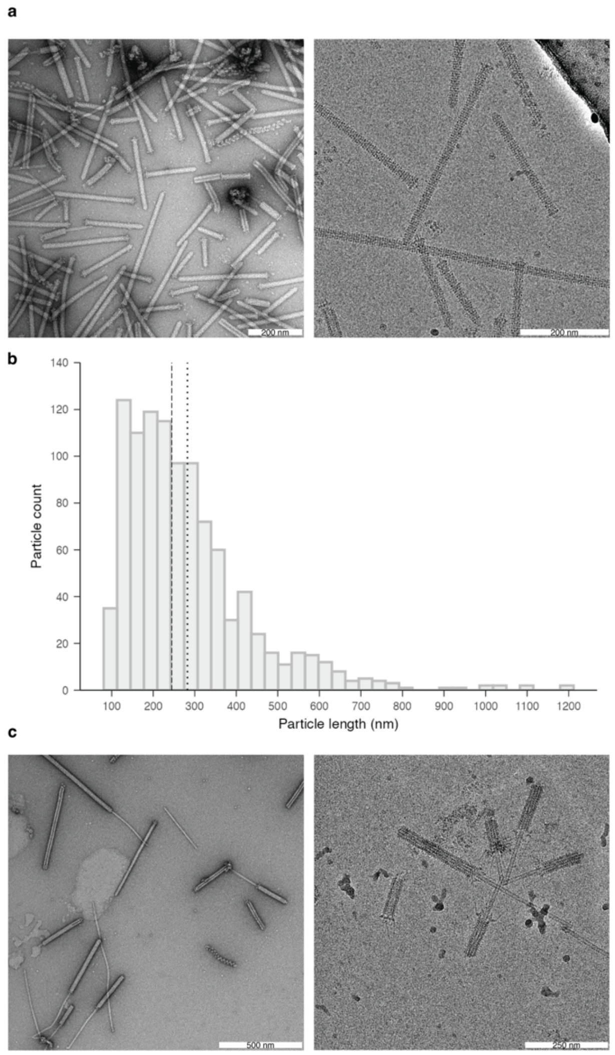


**Supplementary Figure 2. Characterization of the *Pl*PVC1 particle in its extended and contracted states. (a)** Representative NS-EM image (*left*) and micrograph (*right*) displaying *Pl*PVC1 particles in its extended state. **(b)** Length variation of *Pl*PVC1 purified particles. Particle length was measured manually on 1030 particles in NS-EM images. The dashed vertical line indicates the median of the distribution (median = 244.6 nm). The dotted vertical line indicates the mean of the distribution (mean = 282.4 nm). **(c)** Representative NS-EM image (*left*) and micrograph (*right*) displaying *Pl*PVC1 particles in its contracted state.

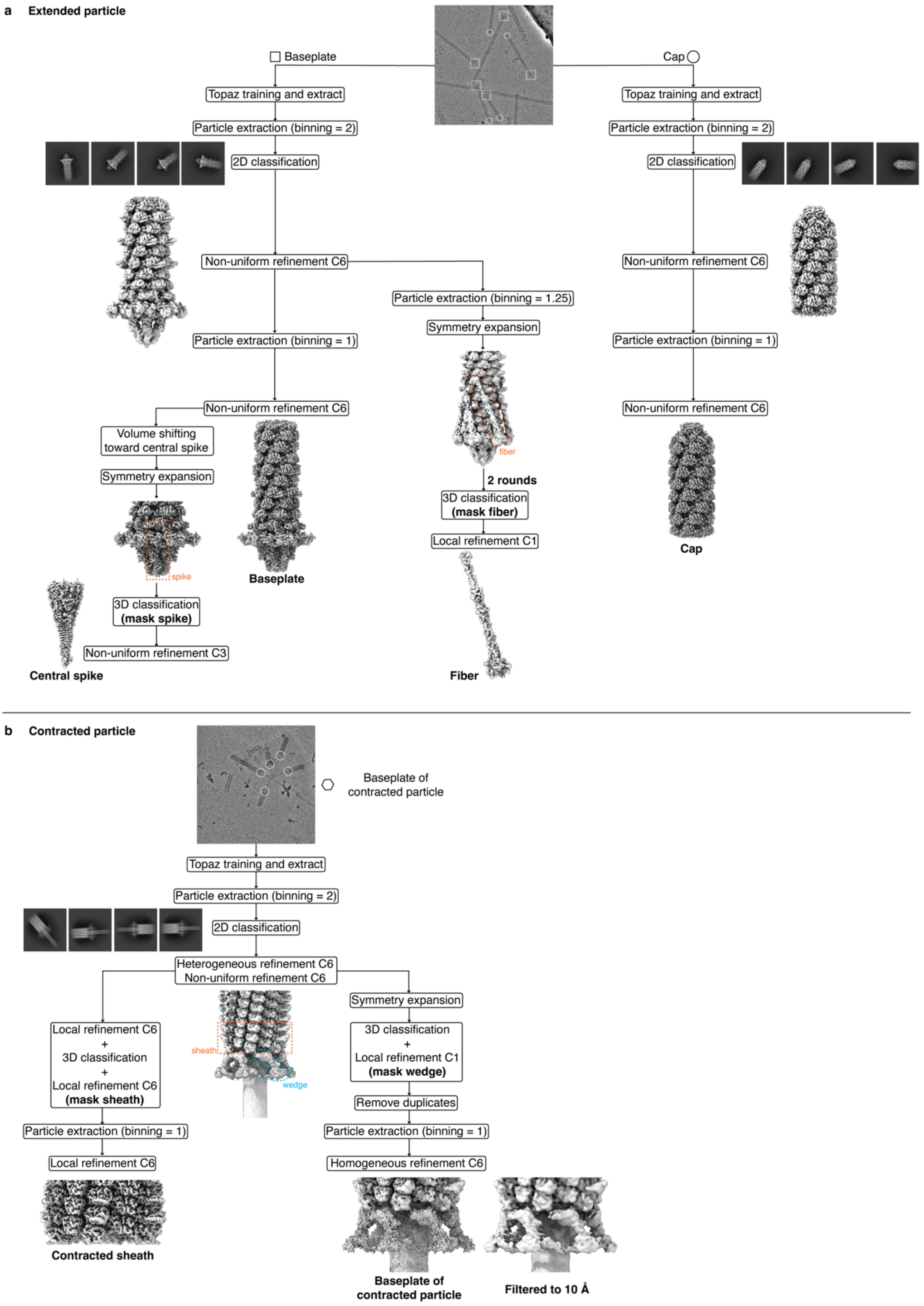


**Supplementary Figure 3. Cryo-EM data processing workflow.** Processing pipeline for the *Pl*PVC1 particle in its extended **(a)** and contracted **(b)** states. Representative micrographs, 2D classes, 3D volumes, and masks are depicted.

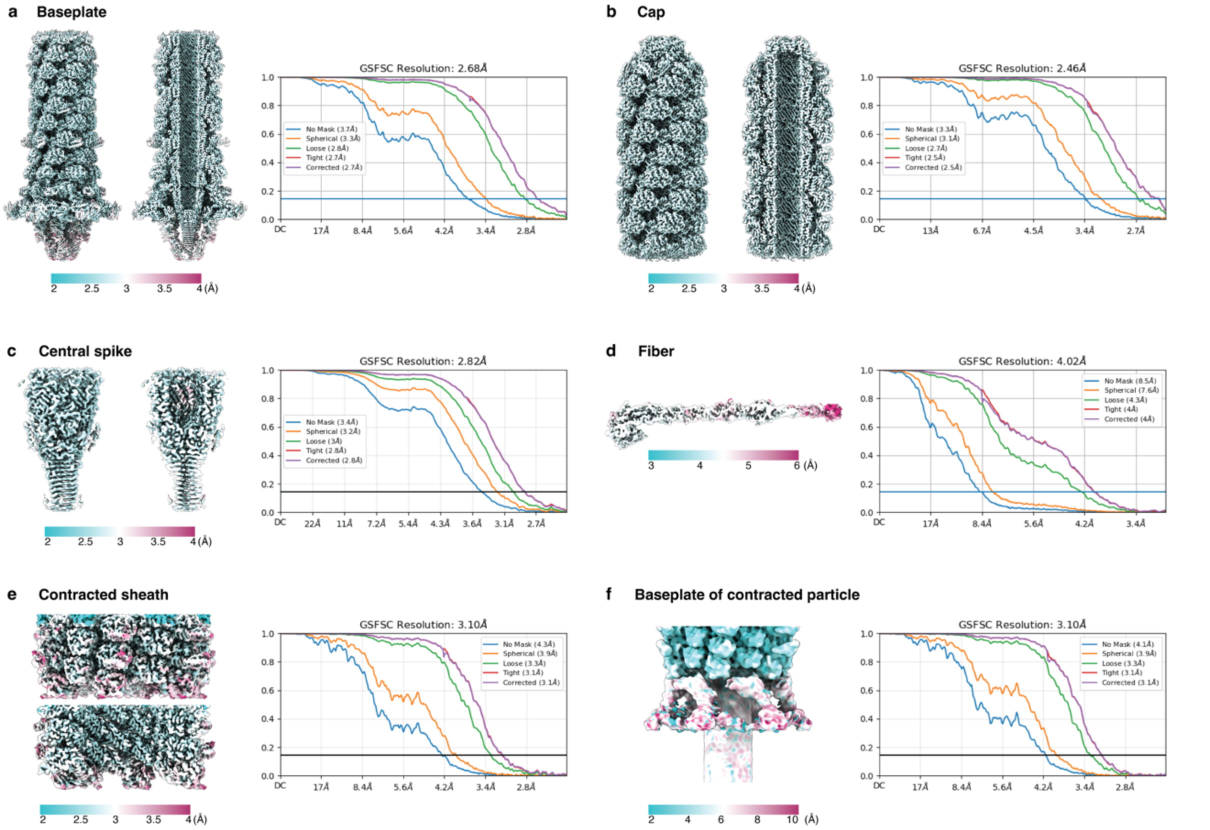


**Supplementary Figure 4. Cryo-EM analysis of reconstructed density maps.** Cryo-EM maps colored by local resolution and gold-standard curves (FSC = 0.143) of **(a)** baseplate, **(b)** cap, **(c)** central spike, **(d)** fiber, **(e)** contracted sheath, and **(f)** baseplate of the contracted particle, filtered to 10 Å.

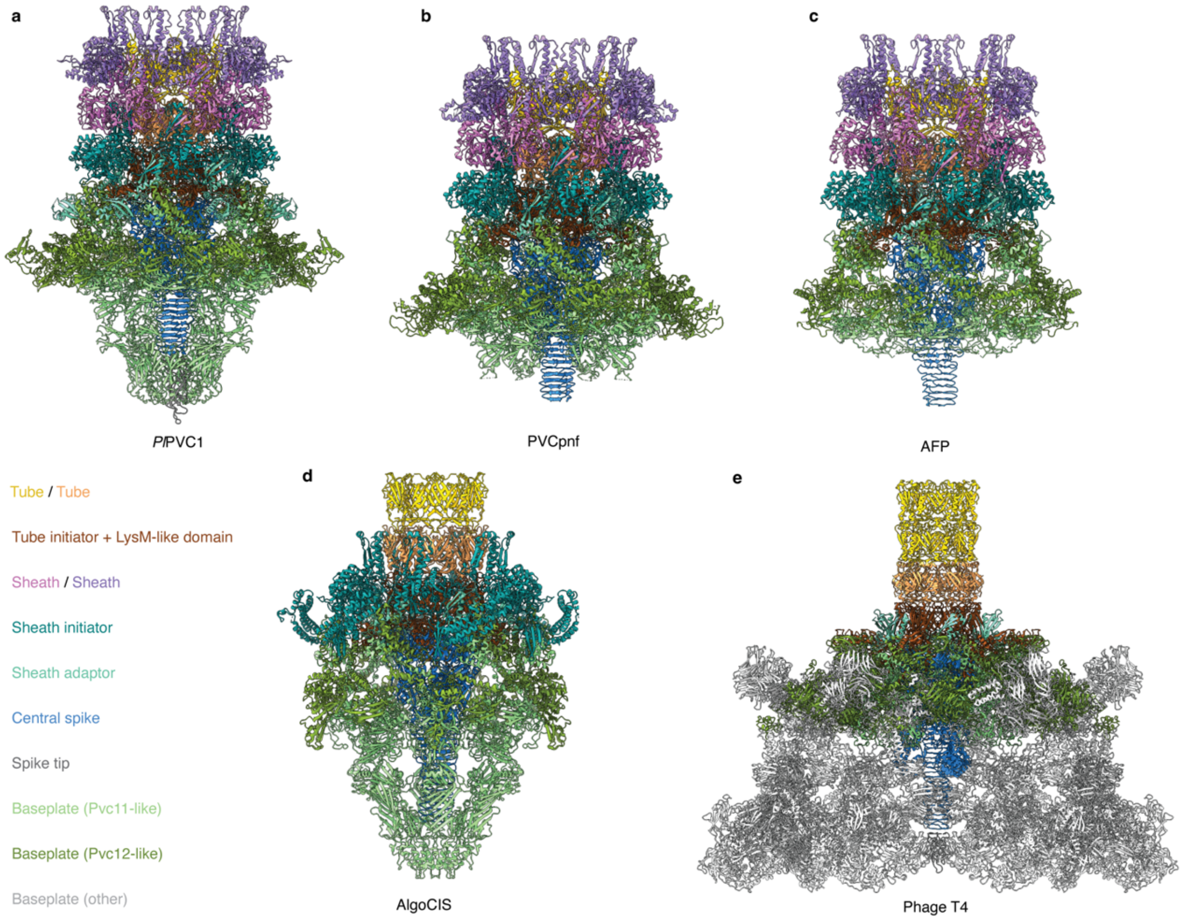


**Supplementary Figure 5. Comparison between baseplate structures of *Pl*PVC1, PVCpnf, AFP, AlgoCIS, and phage T4**. Side views of atomic models of the baseplate complexes in **(a)** *Pl*PVC1, **(b)** PVCpnf [PDBs: 6J0N and 6J0M], **(c)** AFP [PDBs: 6RAO and 6RBK], **(d)** AlgoCIS [PDB: 7AEF], **(e)** phage T4 [PDB: 5IV5]. The different proteins subunits are colored according to **Figure 1** in this manuscript. Tube: gold and light brown. Tube initiator: dark brown. Sheath: purple and pink. Sheath initiator: teal. Sheath adaptor: aquamarine. Central spike: blue. Spike tip: grey. Baseplate wedges: light and dark green.

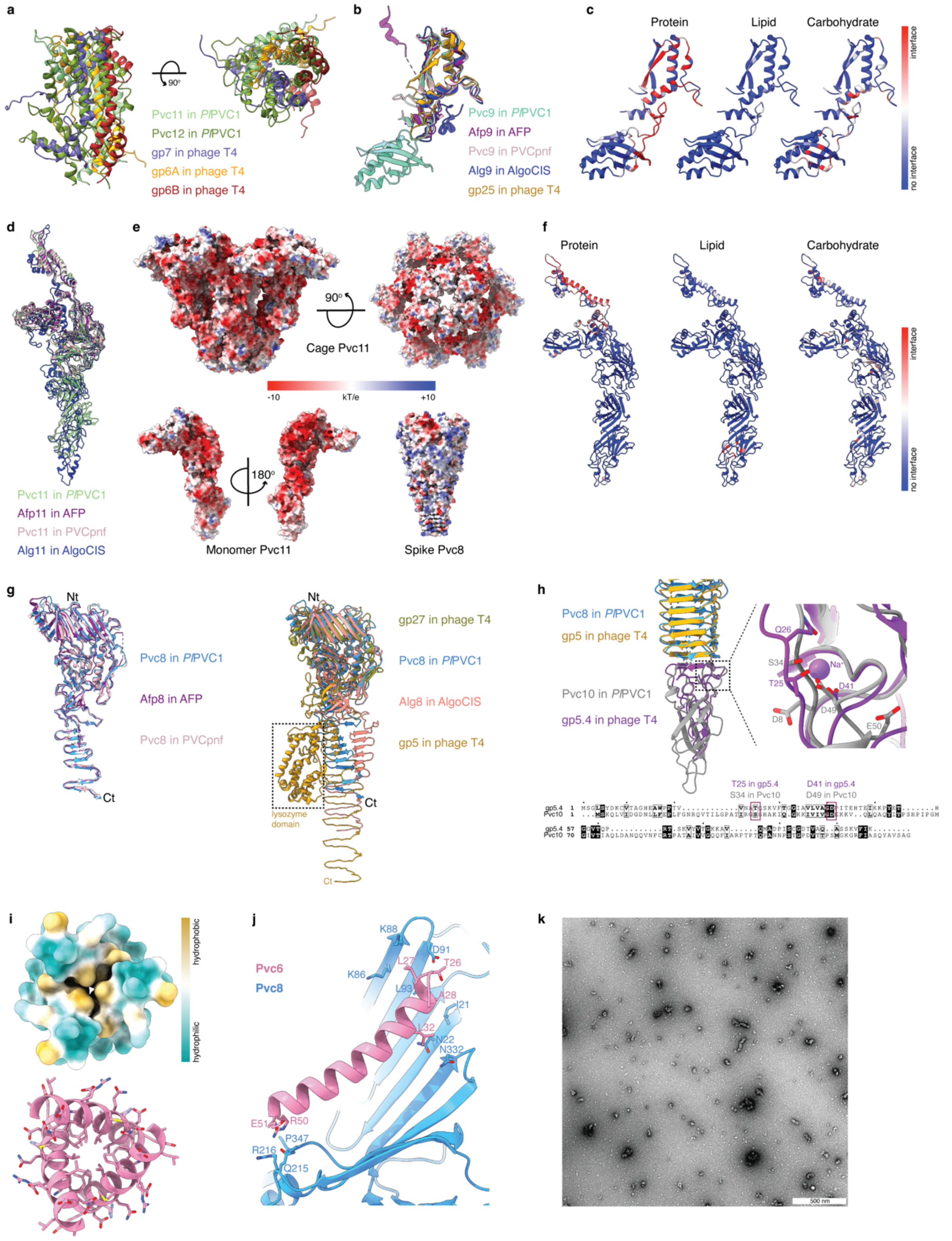


**Supplementary Figure 6. Structural description of the *Pl*PVC1 baseplate, sheath adaptor, cage, spike, tip, and plug. (a)** Close-up comparative views of the helical core bundle of baseplate wedges Pvc11-Pvc12 in *Pl*PVC1 and [gp6]$_2$-gp7 in phage T4. Pvc11 in light green, Pvc12 in dark green, gp7 in blue, gp6A in orange, gp6B in red [PDB: 5IV5, chains u, v, w]. **(b)** Structural comparison of Pvc9, Afp9, Alg9 and gp25. AFP [PDB: 6RAO, chain H], PVCpnf [PDB: 6J0N, chain D], AlgoCIS [PDB: 7AEF, chain S], phage

T4 [PDB: 5IV5, chain T]. **(c)** PeSTo-based prediction of binding interfaces of Pvc9 with proteins, lipids, and carbohydrates. The confidence of the prediction is represented as a gradient, from blue (no interface) to red (interface). **(d)** Structural comparison of Pvc11, Afp11, and Alg11. AFP [PDB: 6RAO, chain I], PVCpnf [PDB: 6J0N, chain J], AlgoCIS [PDB: 7AEF, chain G]. **(e)** Surface representation of *Pl*PVC1 cage, cage monomer, and spike, colored according to electrostatic potential. **(f)** PeSTo-based prediction of binding interfaces of Pvc11 with proteins, lipids, and carbohydrates. The confidence of the prediction is represented as a gradient, from blue (no interface) to red (interface). **(g)** Structural comparison of Pvc8, Afp8, Alg8, gp27, and gp5. AFP [PDB: 6RBK, chain C], PVCpnf [PDB: 6J0M, chain C], AlgoCIS [PDB: 7AEF, chain q], phage T4 [PDB: 5IV5, chains YA and YD]. **(h)** *Left:* Structural comparison of Pvc10 in *Pl*PVC1 and gp5.4 in phage T4 [PDB: 4KU0]. pTM score of Pvc10 AlphaFold model = 0.86. *Right*: Zoom-in view of region involved in $Na^+$ binding, with residues labelled and shown as sticks. *Bottom*: MSA of gp5.4 and Pvc10 sequences, with highlighted conserved residues involved in $Na^+$ binding (conservation calculated with ConSurf Server). **(i)** *Top*: surface representation of Pvc6 colored according to molecular lipophilicity potential, revealing the hydrophobic core in Pvc6. *Bottom*: top view of the atomic model of Pvc6 in ribbon diagram, showing hydrophobic residues in core and charged residues in outer surface. **(j)** Zoom-in view of the main interactions between plug Pvc6 and spike Pvc8. Residues are labeled and shown as sticks. **(k)** NS-EM image showing the absence of assembled *Pl*PVC1 particles in the mutant *Pl*PVC1ΔPvc6 preparation.

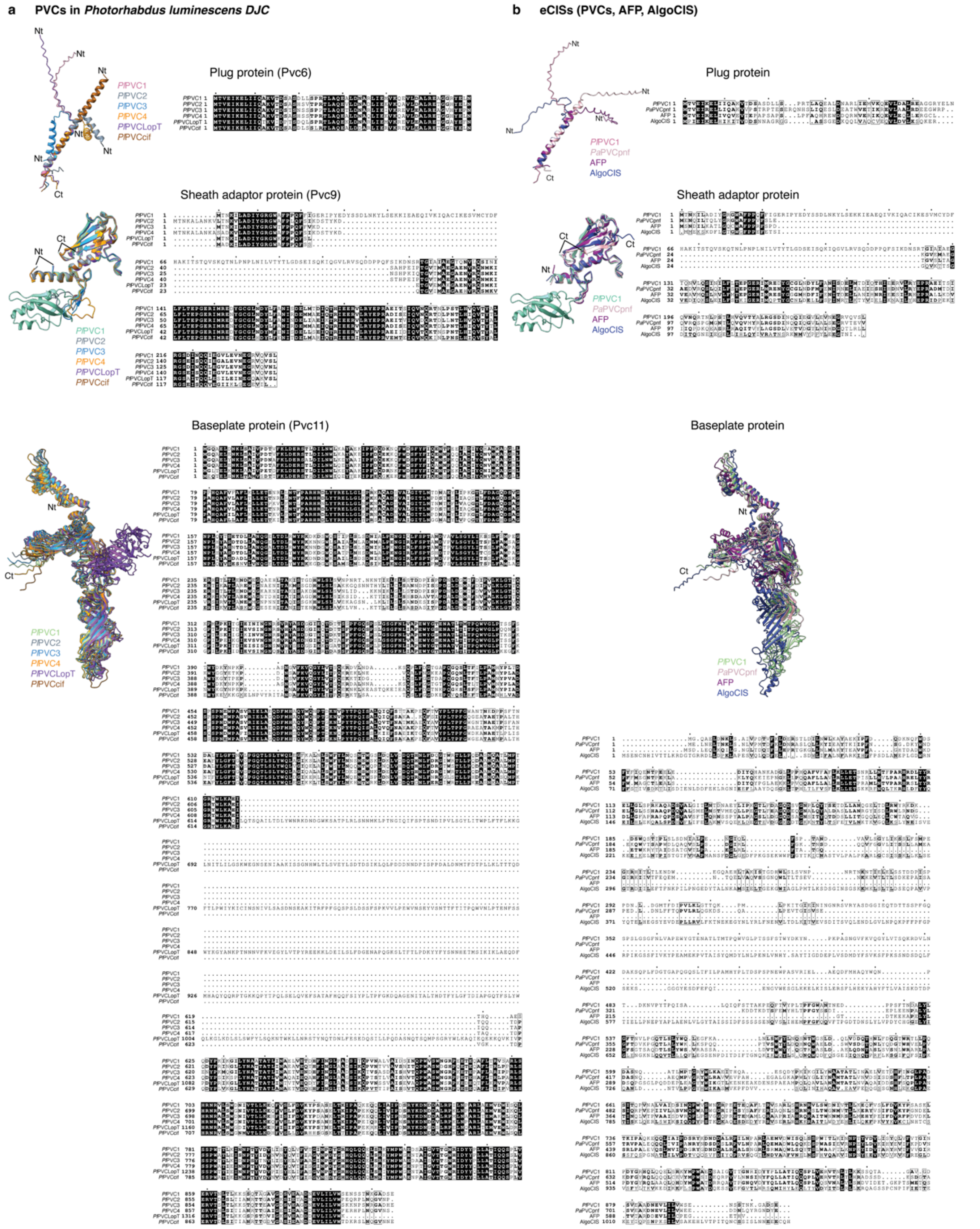


**Supplementary Figure 7. Sequence alignments and structural comparison of the plug proteins, sheath adaptor proteins, and baseplate proteins from PVCs in *P. luminescens DJC* and eCIS. (a)** Structural comparison of AlphaFold predictions and MSA of the sequences of the plug proteins (*top*), sheath adaptor proteins (*middle*), and baseplate proteins (*bottom*) from all the *pvc* operons in *P. luminescens DJC* (*Pl*PVC1, *Pl*PVC2, *Pl*PVC3, *Pl*PVC4, *Pl*PVCLopT, *Pl*PVCcif). The AlphaFold predictions of the plug proteins align between residues 24-51, corresponding with the modeled region of Pvc6 in this study. Nt = N-terminus. Ct = C-terminus. **(b)** Structural comparison of AlphaFold predictions and MSA of the sequences of the plug proteins (*top*), sheath adaptor proteins (*middle*), and baseplate proteins (*bottom*) from *Pl*PVC1, *Pa*PVCpnf, AFP, and AlgoCIS. Nt = N-terminus. Ct = C-terminus.

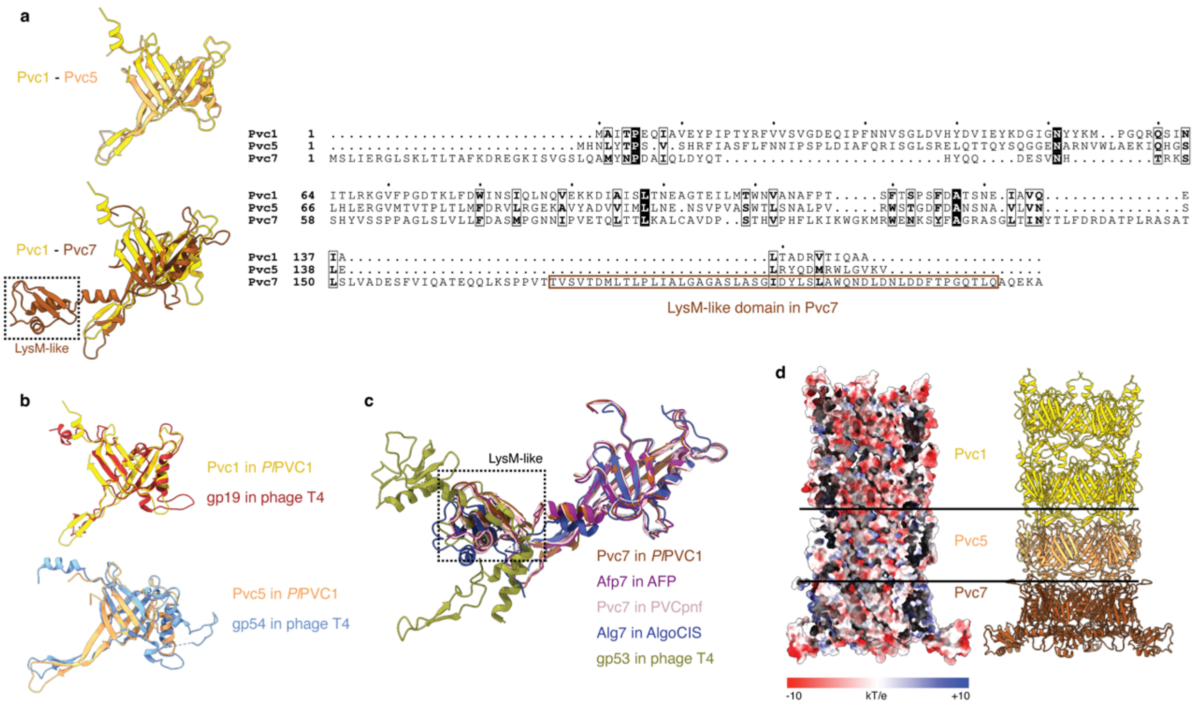


**Supplementary Figure 8. Sequence alignments, structural comparison, and structural analysis of the tube proteins. (a)** *Left*: Structural comparison of tube subunits in *Pl*PVC1, Pvc1-Pvc5 and Pvc1-Pvc7. *Right*: MSA of the sequences of the tube proteins in *Pl*PVC1. The LysM-like domain is marked in both the structure and sequence of Pvc7. **(b)** Structural comparison of Pvc1 and Pvc5 in *Pl*PVC1 and gp19 and gp54 in phage T4 [PDB: 5IV5, chains BB and BG]. **(c)** Structural comparison of Pvc7, Afp7, Alg7, and gp53. The LysM-like domain is marked with a rectangle. AFP [PDB: 6RAO, chain G], PVCpnf [PDB: 6J0N, chain g], AlgoCIS [PDB: 7AEF, chain Q], phage T4 [PDB: 5IV5, chain BF]. **(d)** Cut-out side view of surface representation of *Pl*PVC1 tube, colored according to electrostatic potential, next to ribbon diagram representation of the same area.

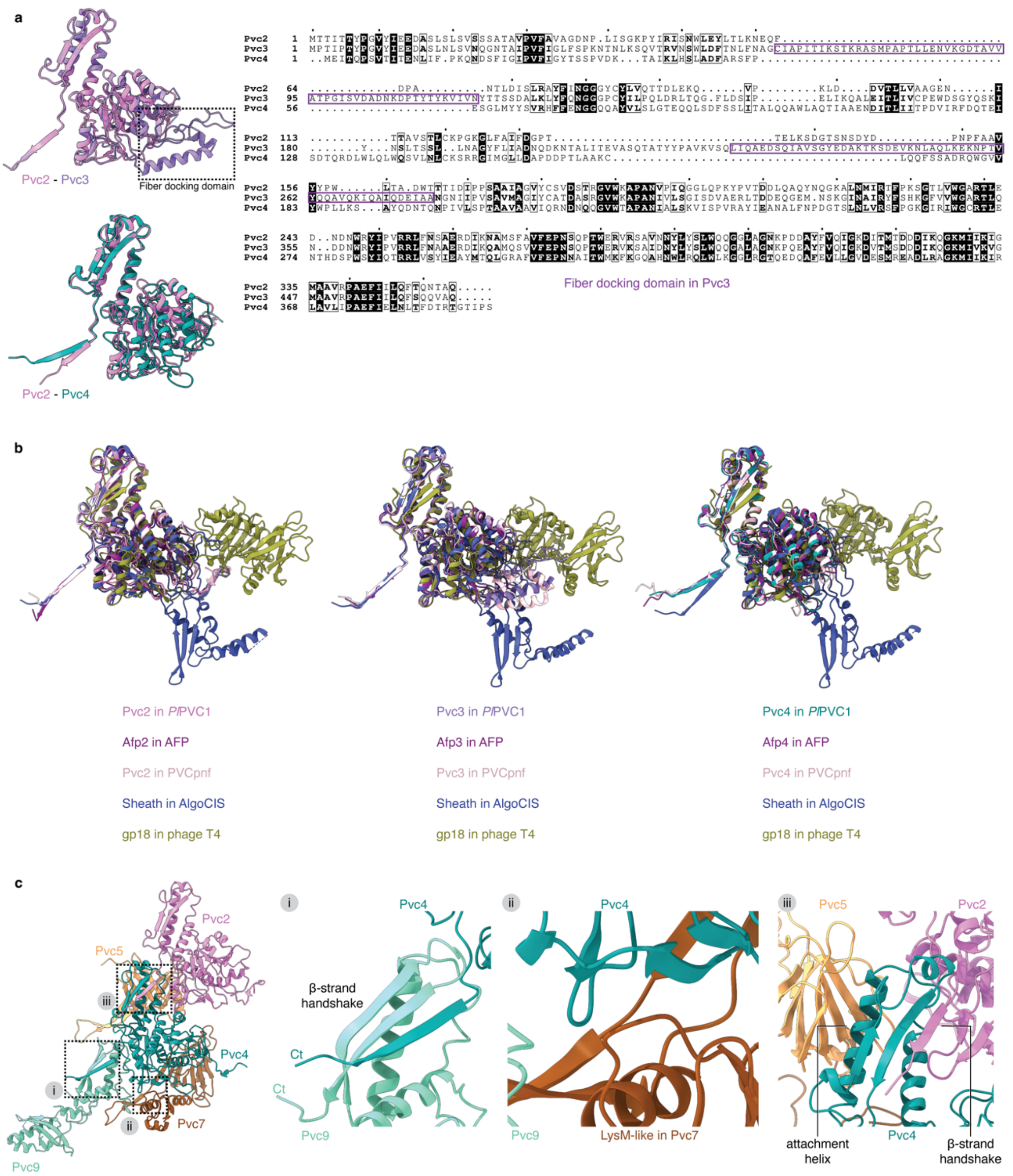


**Supplementary Figure 9. Sequence alignments, structural comparison, and structural analysis of the sheath proteins**. **(a)** *Left*: Structural comparison of sheath subunits in *Pl*PVC1, Pvc2-Pvc3 and Pvc2-Pvc4. *Right*: MSA of the sequences of the sheath proteins in *Pl*PVC1. The fiber docking domain is marked in both the structure and sequence of Pvc3. **(b)** Structural comparison of sheath proteins in several particles. AFP [PDB: 6RAO, chains C, D, E], PVCpnf [PDB: 6J0N, chains m, t, a], AlgoCIS [PDB: 7AE0, chain 5a], phage T4 [PDB: 3J2M, chain U]. **(c)** Interactions between the sheath initiator Pvc4 and sheath and tube proteins. (i) β-strand handshake between Pvc9 and Pvc4. Ct = C-terminus. (ii) Interaction between Pvc4 and LysM-like domain in the tube initiator Pvc7. (iii) β-strand handshake between Pvc4 and Pvc2 in the first layer of the sheath, and attachment helix between Pvc4 and Pvc5 in the first layer of the tube.

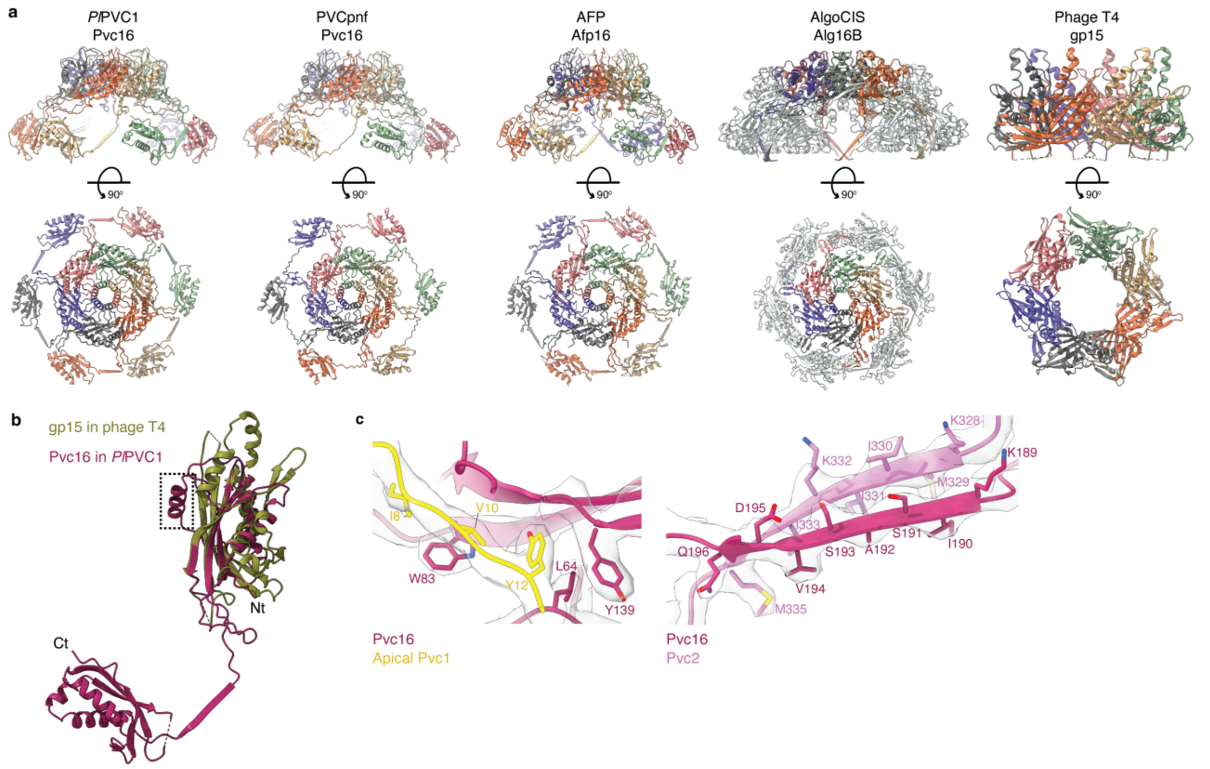


**Supplementary Figure 10. Comparison between cap complexes and structural analysis of the cap in the *Pl*PVC1 particle. (a)** Side and top views of atomic models of the cap complexes in *Pl*PVC1, PVCpnf [PDB: 6J0F], AFP [PDB: 6RAP], AlgoCIS [PDB: 7ADZ], and phage T4 [PDB: 3J2M]. **(b)** Structural comparison of Pvc16 in *Pl*PVC1 and gp15 in phage T4 [PDB: 3J2M, chain A]. The extra α-helix in Pvc16 is marked with a dashed rectangle. Nt = N-terminus. Ct = C-terminus. **(c)** Interactions between Pvc16 and apical Pvc1 subunit (*left*) and interactions in the β-intercalation handshake between Pvc16 and Pvc2 (*right*) in the upmost layer of the particle in its extended state. Residues are labeled and shown as sticks.

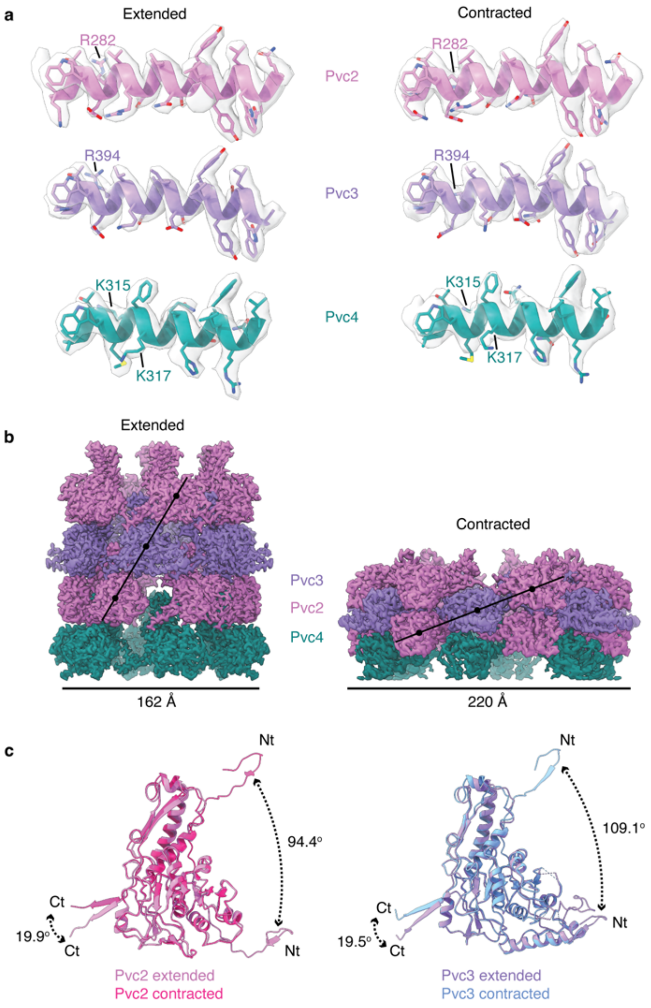


**Supplementary Figure 11. Structural comparison of sheath proteins in the extended and contracted states. (a)** Cryo-EM density and ribbon diagram for the tube-sheath attachment helices in sheath proteins, before (*left*) and after (*right*) treatment with 3 M urea for contraction. Pink helix in Pvc2, from residues 277 to 297. Purple helix in Pvc3, from residues 390 to 408. Teal helix in Pvc4, from residues 310 to 327. Slight changes can be appreciated in R282 in Pvc2, R394 in Pvc3, and K315 and K317 in Pvc4. The core structure of the helices remains intact after treatment with 3 M urea. Residues are labeled and shown as sticks. **(b)** Side views of cryo-EM maps of extended (*left*) and contracted (*right*) sheath, with diameter measurements. **(c)** Structural comparison of sheath proteins Pvc2 and Pvc3 in the extended and contracted states. Rearrangement of both C- and N-termini can be seen. Nt = N-terminus. Ct = C-terminus.

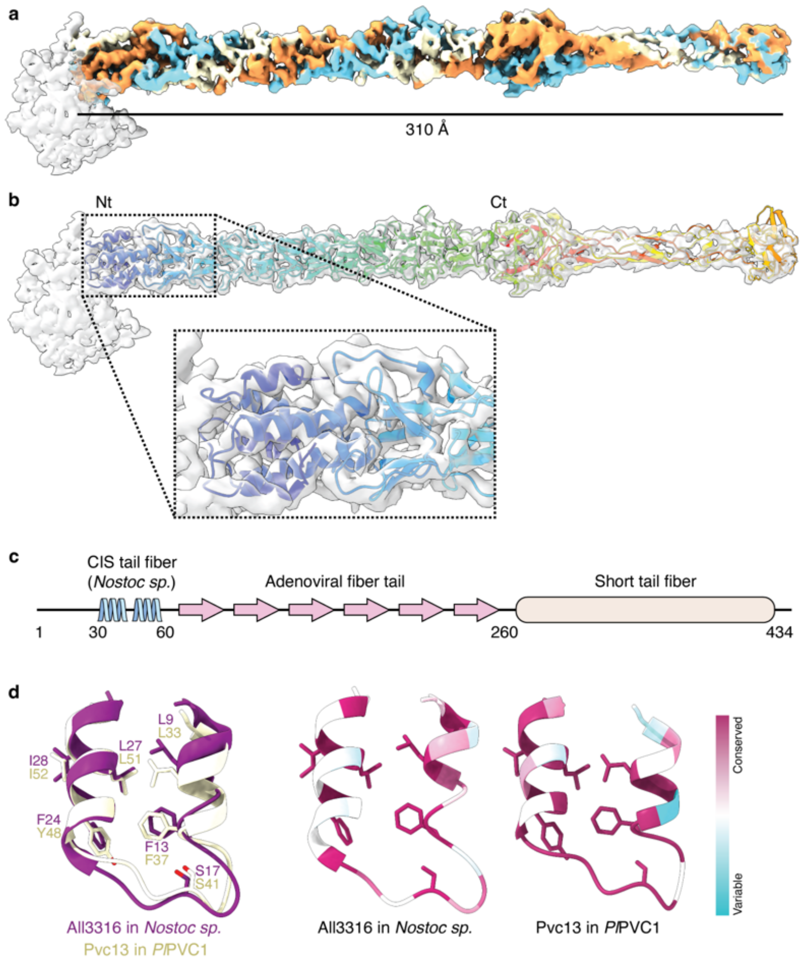


**Supplementary Figure 12. Molecular organization of the *Pl*PVC1 fiber. (a)** Cryo-EM density of the fiber showing intertwined copies of Pvc13 trimer. Each chain of Pvc13 is colored in cyan, yellow, and orange, respectively. **(b)** *Top:* Arched topology of the *Pl*PVC1 fiber. Cryo-EM density and ribbon diagram of the fiber, colored in rainbow palette, where blue corresponds to N-terminal region and red to C-terminal region. The AlphaFold model of the fiber (pTM score = 0.68) was fitted in the cryo-EM density of the fiber. Nt = N-terminus. Ct = C-terminus. *Bottom*: Zoom-in view of the helices in the N-terminal region of the fiber. **(c)** Schematic representation of the Pvc13 sequence organization. The N-terminal region shows homology to CIS fibers. The middle region shows homology to adenoviral fibers. The C-terminal region shows homology to host-binding domains of short tail fibers from bacteriophages. Regions were depicted based on statistically significant hits in HHpred (>95%). **(d)** *Left*: Structural comparison of conserved helices in All3316 from *Nostoc sp.* (strain PCC 7120 / SAG 25.82 / UTEX 2576) [PDB: 7B5H, chain AE] and helices in Pvc13 in *Pl*PVC1. Conserved matching residues are labeled and shown as sticks. *Right*: Conservation (calculated with ConSurf Server) of residues 5-31 in All3316 and 29-55 in Pvc13.

**Supplementary Table 1: *Pl*PVC1 proteins and their homologs in other contractile systems.**

| PVC | AFP | Phage T4 | T6SS | Pyocin R2 | AlgoCIS | Function |
|---|---|---|---|---|---|---|
| Pvc1 | Afp1 | gp19 | Hcp | PA0623 | Alg1 | Tube |
| Pvc2-3-4 | Afp2-3-4 | gp18 | TssB/C | PA0622 | Alg2 | Sheath |
| Pvc5 | Afp5 | gp54 | Hcp | PA0623 | Alg5 | Tube (second layer) |
| Pvc6 | Afp6 | - | - | - | Alg6 | Plug |
| Pvc7 | Afp7 | gp48/gp53 LysM | Hcp | PA0623 / PA0627 LysM | Alg7 | Tube initiator (first layer) |
| Pvc8 | Afp8 | gp27/gp5 | VgrG | PA0616 / PA0628 | Alg8 | Central spike |
| Pvc9 | Afp9 | gp25 | TssE | PA0617 | Alg9 | Sheath adaptor |
| Pvc10 | Afp10 | gp5.4 | VgrG PAAR | PA0616 | Alg10 | Spike tip |
| Pvc11 | Afp11 | gp6 | TssF | PA0618 | Alg11 | Baseplate wedge |
| Pvc12 | Afp12 | gp6/gp7 | TssF/G | PA0619 | Alg12 | Baseplate wedge |
| Pvc13 | Afp13 | gp12 | - | PA0620 | - | Tail fiber |
| Pvc14 | Afp14 | gp29 | - | PA0625 | Alg14 | Tape measure protein |
| Pvc15 | Afp15 | - | ClpV | - | Alg15 | AAA+ ATPase |
| Pvc16 | Afp16 | gp15 | TssA | PA0626 / PA0615 | Alg16A | Cap / Tube-sheath terminator |

**Supplementary Table 2: Particle length statistics**.

| | |
|---|---|
| Particle count | 1030 |
| Minimum length | 89.48 nm |
| Maximum length | 1188.18 nm |
| Mean | 282.4 nm |
| Median | 244.6 nm |
| Standard deviation | 154.9 nm |

**Supplementary Table 3. Cryo-EM data collection, refinement, and validation statistics.**

| **Data collection and processing** | **Baseplate (EMD-53137)** | **Central spike (EMD-53138)** | **Fiber (EMD-53140)** | **Cap (EMD-53139)** | **Contracted sheath (EMD-53141)** | **Baseplate contracted particle (EMD-53143)** |
|---|---|---|---|---|---|---|
| Microscope | Titan Krios G2 | Titan Krios G2 | Titan Krios G2 | Titan Krios G2 | Titan Krios G2 | Titan Krios G2 |
| Magnification (nominal) | 105,000x | 105,000x | 105,000x | 105,000x | 105,000x | 105,000x |
| Voltage (kV) | 300 | 300 | 300 | 300 | 300 | 300 |
| Electron exposure ($e^-/Å^2$) | 41 | 41 | 41 | 41 | 40 | 40 |
| Defocus range ($\mu$m) | -2 to -0.6 | -2 to -0.6 | -2 to -0.6 | -2 to -0.6 | -2 to -0.6 | -2 to -0.6 |
| Pixel size (Å) | 1.2 | 1.2 | 1.2 | 1.2 | 1.2 | 1.2 |
| Micrographs used (no.) | 17445 | 17445 | 17445 | 17725 | 11896 | 11702 |
| Final particles (no.) | 53537 | 47836 | 71148 (sym.exp.) | 57353 | 30862 | 28687 |
| Box size (pixels) | 700 | 360 | 560 | 560 | 700 | 700 |
| Symmetry imposed | C6 | C3 | C1 | C6 | C6 | C6 |
| Map sharpening B-factor ($Å^2$) | 56.8 | 59.1 | 50.3 | 48.7 | 73.9 | 68 |
| Map resolution (Å) (FSC 0.143) | 2.68 | 2.82 | Range 4-6 | 2.46 | 3.10 | Range 4-10 |

| **Refinement and validation** | **Baseplate (9QGL)** | **Central spike (9QGM)** | **Baseplate-Fiber (9QGO)** | **Cap (9QGN)** | **Contracted sheath (9QGP)** |
|---|---|---|---|---|---|
| Model composition | | | | | |
| Chains | 54 | 6 | 6 | 24 | 24 |
| Non-hydrogen atoms | 171084 | 12963 | 1505 | 43476 | 67578 |
| Protein residues | 21576 | 1665 | 193 | 5562 | 8646 |
| Ligands | 0 | 0 | 0 | 0 | 0 |
| B-factors (mean; $Å^2$) | | | | | |
| Protein | 87.70 | 80.19 | 121.57 | 81.09 | 87.65 |
| Ligands | - | - | - | - | - |
| R.m.s deviations | | | | | |
| Bond lengths (Å) | 0.002 | 0.003 | 0.004 | 0.002 | 0.002 |
| Bond angles (°) | 0.465 | 0.531 | 0.866 | 0.424 | 0.492 |
| CC (mask) | 0.90 | 0.88 | 0.78 | 0.91 | 0.89 |
| MolProbity score | 1.41 | 1.41 | 2.02 | 1.22 | 1.66 |
| Poor rotamer (%) | 1.61 | 1.44 | 0.60 | 1.05 | 2.08 |
| Ramachandran plot | | | | | |
| Favored (%) | 98.01 | 98.00 | 97.21 | 98.20 | 97.76 |
| Allowed (%) | 1.99 | 2.00 | 2.79 | 1.80 | 2.24 |
| Disallowed (%) | 0.00 | 0.00 | 0.00 | 0.00 | 0.00 |

**Supplementary Table 4: Percentage of identity between proteins in *Pl*PVC1 and proteins in other PVCs from *P. luminescens DJC*.**

| ***Pl*PVC1** | **Tube 1** | **Sheath 2** | **Sheath 3** | **Sheath 4** | **Tube 5** | **Plug 6** | **Tube 7** | **Spike 8** | **Sheath 9** | **Tip 10** | **Baseplate 11** | **Baseplate 12** | **Fiber 13** | **tmp 14** | **ATPase 15** | **Cap 16** |
|---|---|---|---|---|---|---|---|---|---|---|---|---|---|---|---|---|
| *Pl*PVC2 | 86,58 | 74,72 | 71,21 | 74,62 | 91,45 | 83,05 | 87,67 | 81,61 | 72,55 | 89,86 | 75,31 | 75,05 | 51,08 | 70,82 | 81,1 | 78 |
| *Pl*PVC3 | 91,28 | 79,1 | 74,89 | 84,91 | 92,76 | 79,66 | 83,7 | 78,24 | 78,38 | 80,43 | 76 | 79,79 | 48,15 | 65,82 | 83,86 | 83,48 |
| *Pl*PVC4 | 88,59 | 74,01 | 74,02 | 75,13 | 92,11 | 84,75 | 88,11 | 81,24 | 72,55 | 88,41 | 75,83 | 73,19 | 57,29 | 74,36 | 82,42 | 82,67 |
| *Pl*PVCLopT | 87,92 | 75,42 | 56,37 | 74,35 | 90,79 | 81,36 | 85,46 | 81,8 | 75 | 84,06 | 70,42 | 65,6 | N/A | 69,05 | 77,78 | 70 |
| *Pl*PVCcif | 67,79 | 77,12 | 73,26 | 73,66 | 94,74 | 86,44 | 82,38 | 79,55 | 69,57 | 88,41 | 68,92 | 70,75 | 55,39 | 58,03 | 78,24 | 66,89 |

**Supplementary Table 5: Percentage of identity between proteins in *Pl*PVC1 and proteins in other eCIS.**

| ***Pl*PVC1** | **Tube 1** | **Sheath 2** | **Sheath 3** | **Sheath 4** | **Tube 5** | **Plug 6** | **Tube 7** | **Spike 8** | **Sheath 9** | **Tip 10** | **Baseplate 11** | **Baseplate 12** | **Fiber 13** | **tmp 14** | **ATPase 15** | **Cap 16** |
|---|---|---|---|---|---|---|---|---|---|---|---|---|---|---|---|---|
| *Pa*PVCpnf | 79,19 | 70,17 | 51,04 | 53,96 | 80,92 | 49,12 | 59,47 | 44,36 | 60 | 53,62 | 49,1 | 53,62 | 41,77 | 65,26 | 56,98 | 52,22 |
| AFP | 73,83 | 55,3 | 49,1 | 44,76 | 69,8 | 43,64 | 62,56 | 36,95 | 43,57 | 48,18 | 39,83 | 44,57 | 30,88 | 30,18 | 53,35 | 45,42 |
| AlgoCIS | 40,29 | 40,24 | 40,72 | 31,55 | 29,93 | 35,29 | 24,54 | 25,35 | 29,2 | 19,57 | 24,37 | 22,04 | N/A | 19,96 | 32,49 | 18,52 |
